# Three-dimensional spike localization and improved motion correction for Neuropixels recordings

**DOI:** 10.1101/2021.11.05.467503

**Authors:** Julien Boussard, Erdem Varol, Hyun Dong Lee, Nishchal Dethe, Liam Paninski

## Abstract

Neuropixels (NP) probes are dense linear multi-electrode arrays that have rapidly become essential tools for studying the electrophysiology of large neural populations. Unfortunately, a number of challenges remain in analyzing the large datasets output by these probes. Here we introduce several new methods for extracting useful spiking information from NP probes. First, we use a simple point neuron model, together with a neural-network denoiser, to efficiently map single spikes detected on the probe into three-dimensional localizations. Previous methods localized individual spikes in two dimensions only; we show that the new localization approach is significantly more robust and provides an improved feature set for clustering spikes according to neural identity (“spike sorting”). Next, we denoise the resulting three-dimensional point-cloud representation of the data, and show that the resulting 3D images can be accurately registered over time, leading to improved tracking of time-varying neural activity over the probe, and in turn, crisper estimates of neural clusters over time. Open source code is available at https://github.com/int-brain-lab/spikes_localization_registration.git.

## 1 Introduction

Neuropixels (NP) probes are dense linear multi-electrode arrays that enable the simultaneous observation of hundreds of neurons across multiple brain areas. Since their introduction in [13] (see also [27]), they have rapidly become an essential neurotechnology, deployed in hundreds of labs around the world. A number of challenges remain in analyzing the large datasets output by these probes. The basic goal is “spike sorting” — i.e., to detect action potentials (“spikes”) and assign these spikes to individual neurons. Despite significant effort in the field [4, 12, 21, 27], current spike sorters for NP data have trouble tracking temporally non-stationary data and accurately sorting small spikes [28].

One major advantage of NP probes is their spatial density: the high spatial resolution of electrodes on the probe implies that each extracellular spike will typically be detected at multiple sites, providing an opportunity to “triangulate” the location of each spike. If effective, this spike localization can be helpful for multiple downstream tasks: spike locations serve as useful low-dimensional summarizations of the high-dimensional spatiotemporal spiking signals, which can in turn be visualized and used as features for clustering and tracking as the probe undergoes small motion relative to the brain.

Several spike sorting algorithms localize spikes using a simple center of mass (CoM) method. More precisely, for each waveform, they select a set of channels, and compute the weighted average of the selected channels’ positions, where the weights are given by an estimate of the amplitude of the spike on each channel. This fast, cheap method gives informative location estimates for spikes in multi-electrode array (MEA) recordings where long dendritic or axonal signals are not prevalent [23, 20, 10] (cf. the primate retina [17], where long axonal signals make localization approaches less useful). However, by definition, the CoM approach localizes spikes inside the convex hull of the electrodes, which is especially problematic for Neuropixels recordings, due to the long and thin shape of the electrodes. Hurwitz et al. [11] propose an improved localization method that provides location estimates that can lie outside of the probe, but the estimates are still (like the CoM method) restricted to the two-dimensional plane spanned by the electrode locations. In addition, several approaches have been developed for localizing *templates* (averages over many spikes, post spike-sorting) in three dimensions, using biophysical models of varying sophistication [26, 25, 8, 5]. Finally, triangulation approaches based on point-source [6, 15] or dipole models [19] have been proposed for smaller-scale tetrode recordings, in which spikes are only detected on four electrodes, limiting the resolution of the resulting localization in the case of small, noisy spiking units.

In this paper, we build on this previous work, making four key contributions:

1. We introduce a denoise-then-triangulate approach, based on a simple point neuron model, that localizes single spikes in three dimensions.
2. This improved three-dimensional localization in turn enables better spatial clustering of distinct neural units characterized by unique action potential waveform shapes.
3. Our localization further enables better motion estimation in recordings with drift.
4. Our approach is robust to different probe geometries. We demonstrate its utility and performance in both Neuropixels 1.0 and 2.0 probes.

## 2 Methods

### 2.1 Point neuron model

A neuron’s spike is the result of a quick change in its membrane potential due to the shift of its charged particles from one side of the membrane to the other. This movement of charge creates an electric field outside of the neuron [14], and Neuropixels electrodes record the potential of this field, at multiple locations along the probe (**Fig. 1A**). We use a simple point-source model for these voltage differences at distance *r* from the cell:

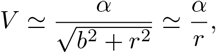

where *α* represents the cell’s overall signal magnitude, and the parameter *b* is fixed independently of the cell. This point-source model has been previously applied in tetrode recordings [6, 15], and could be replaced with more detailed models (dipole, ball-and-stick, etc) if desired.

**Figure 1:**
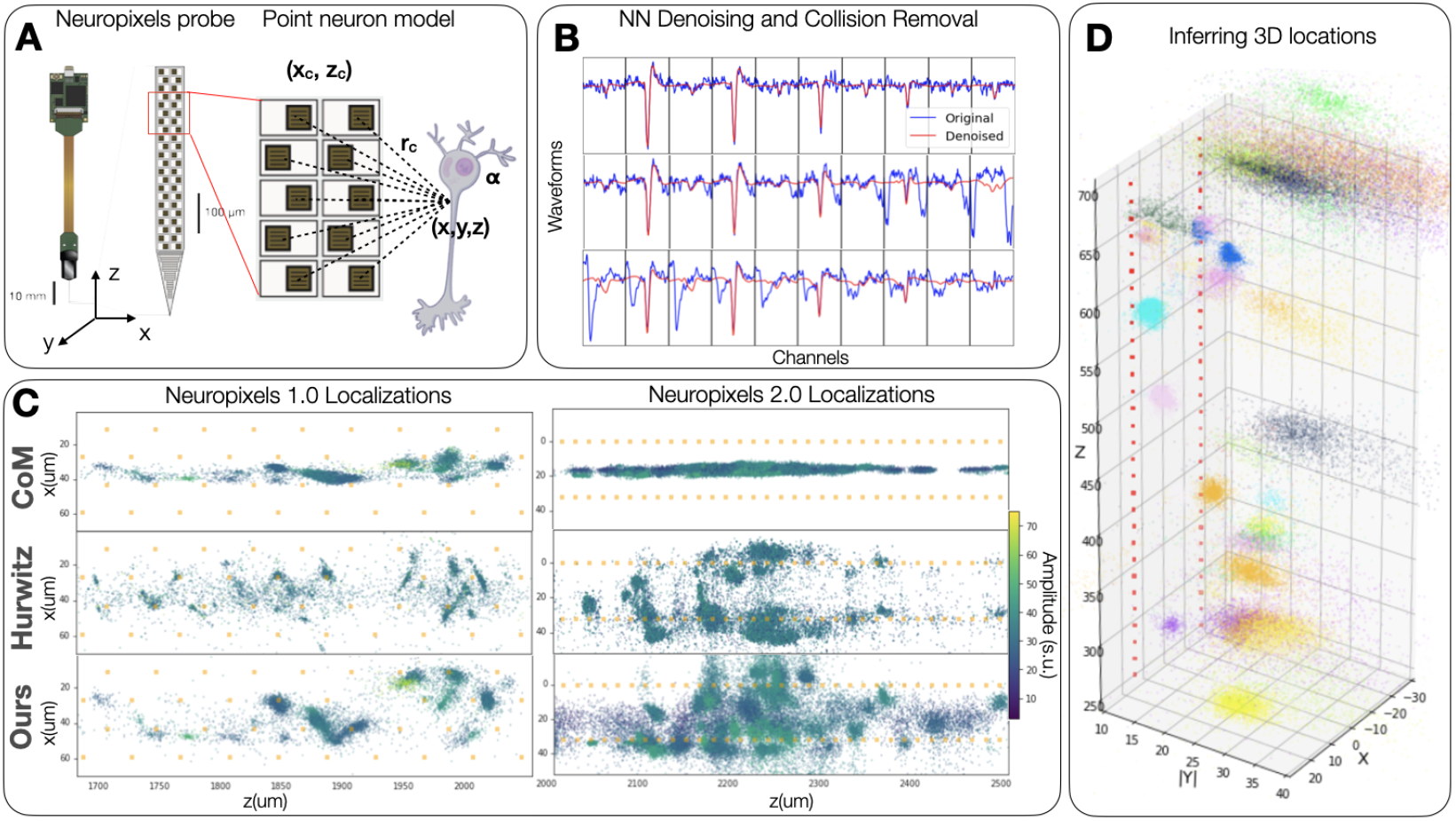
Overview of the proposed localization technique. (**A**) The peak-to-peak voltage amplitude attenuation recorded at each channel of the Neuropixels probe is modelled as approximately inversely proportional to the distance to a spiking unit point source. The 3D spatial location of the spiking unit is then inferred by essentially solving a triangulation problem (minimizing a least-squares loss to all the channels that detected the spiking event). (**B**) Since the triangulation depends on the peak-to-peak amplitudes of spiking events, we denoise the amplitudes using a neural net and further disambiguate distinct units that have spiked at the same time (collision removal). Each row shows a single spike event (original and denoised), captured on multiple nearby electrodes. (**C**) We apply our method to spatially map the localizations of over one million spiking events in both Neuropixels 1.0 and Neuropixels 2.0 recordings. Our localization is compared with the state-of-the-art localization method proposed in [11], as well as the baseline method of inferring position using center of mass (CoM) of channel positions weighted by the recorded amplitude. Spikes are colored by maximum amplitude, normalized by the recording baseline noise level (‘standardized units’ / ‘s.u.’). Yellow corresponds to high and dark blue to low amplitude. (**D**) Our localization yields groups of spiking events that disperse in 3D space and cluster based on amplitude. Colors correspond to clusters of waveforms found by YASS [17], for visualization purposes only. Orange squares represent recording electrode positions. Datasets from https://github.com/flatironinstitute/neuropixels-data-sep-2020/blob/master/doc/cortexlab1.md [27]; see video-figure-1.mp4 for a Datoviz [24] visualization.

### 2.2 Spike denoising and collision suppression

The triangulation approach described below relies only on the amplitude of the spike on each electrode. In practice we measure this amplitude using the peak-to-peak (PTP) value (maximum minus minimum voltage over a short temporal interval). In the presence of noise, the PTP is biased upwards (since noise will increase the maximum and decrease the minimum signal), and this bias depends on the underlying signal strength (the PTP of small spikes is more biased than large spikes). Furthermore, collisions from near-synchronous spike events are prevalent in dense multi-electrode data [17], and even small collisions can lead to inaccurate location estimates if not accounted for properly.

Therefore, it is critical to incorporate a denoising / collision-suppression step prior to triangulation. We use the neural net denoiser implemented in YASS [17] (retrained on NP data), which takes advantage of the waveform’s spatiotemporal signature, to suppress both noise and collisions (Note that simple image denoising methods such as [16, 7, 3] do not serve to suppress collisions, and are therefore less effective here.). **Fig. 1B** shows three examples of waveforms (in blue) and their denoised version (in red). The neural net denoiser removes noise and collided spikes, allowing us to successfully estimate each spike’s PTP amplitude; see the Appendix for further details.

### 2.3 Spike localization

Given the denoised PTP amplitudes obtained above, we can proceed with localizing the origin of each waveform with respect to the multiple electrodes on the NP probe. We will denote {*x, y, z*} the coordinates of the point neuron, and {*x*_*c*_, *z*_*c*_} the planar coordinates of each NP electrode, with *y*_*c*_ set to zero by convention. Here we exploit the fact that each spike is detectable on multiple channels simultaneously: i.e., we have multiple observations of the form 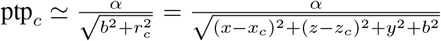, and our goal is to infer *x, y, z*, and *α*. Thus, for each spike and selected set of channels 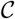 around the “main” channel for each spike (i.e., the channel with the largest PTP amplitude), we want to optimize:

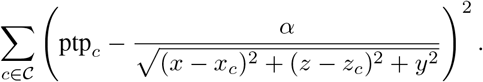

See Appendix for full optimization details. (One important note here: due to the planar NP electrode geometry, the sign of *y* is not recoverable, i.e., we can’t infer whether a spike is in front of or behind the probe, even if we can infer the orthogonal distance from the probe plane.) **Fig. 1 (C)** shows examples of inferred {x,z} locations of detected spikes in both NP1.0 and 2.0 probes (which have different electrode layouts), colored by max PTP. Panel (D) shows an example of 3-d {x, y, z} localization for the NP2.0 data.

### 2.4 Point cloud to image denoising

The estimated localizations are a point cloud in continuous space (**Fig. 2A**). To enable the use of image registration techniques, we convert the point cloud representation to an image representation by generating spatial histograms where pixel location and intensity capture the spatial coordinates of spikes and their mean amplitudes (**Fig. 2B**), following [27]. Note that this image representation is sparse and resembles a speckled Poisson image [18], due to the limited number of spikes that are captured in each spatial position. In order to compute more accurate motion estimates, it is crucial to appropriately denoise this image representation. We adopt a Poisson denoising technique [18], with three steps. First, we apply the Anscombe root transformation [1] to the histogram. Second, we use a Gaussian denoiser (e.g. Block-matching and 3D filtering [7], or non-local means [3]) on the transformed histogram image. Third, the denoised signal is obtained by applying an inverse transformation to the denoised transformed histogram. **Fig. 2C** demonstrates the output for one second of time-binned data: Poisson denoising yields a more continuous pixelwise representation of the spatial histograms, which satisfies the pixel-wise continuity assumptions of image registration algorithms [9] and enables a more robust estimate of displacement.

**Figure 2:**
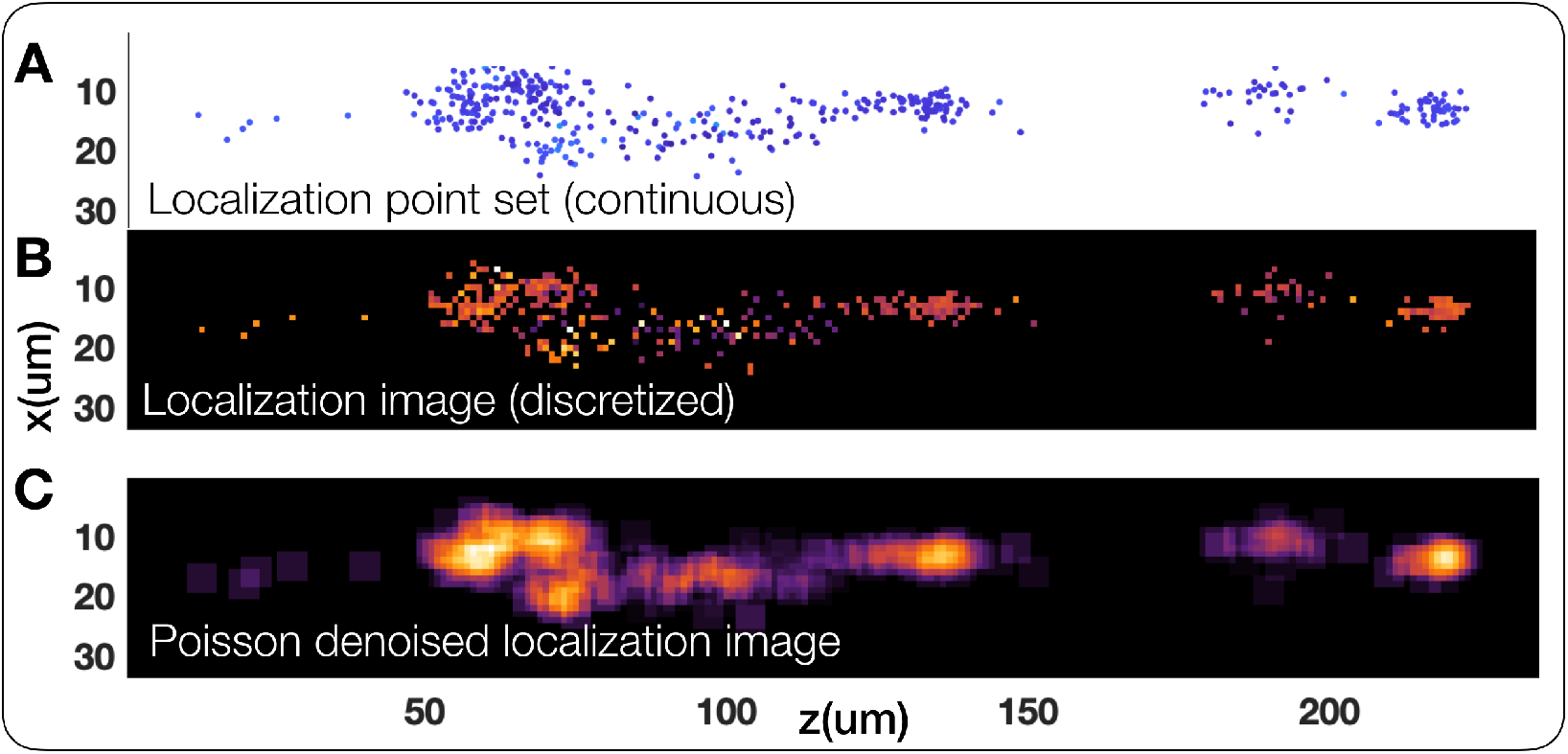
Poisson denoising point sets of localizations yields interpretable images of spiking unit locations. (**A**) The estimated spiking unit localizations are a sparse point set in continuous space. (**B**) By binning spike locations in a spatial histogram, we generate a localization “image" from the point set input, where pixel intensities that denote the average amplitude in positions in space roughly follow a Poisson distribution. Note that this image is also sparse. (**C**) Modelling the noise distribution of this image as Poissonian enables the use of Poisson image denoising techniques [18]. The resulting de-sparsified pixelwise representation of spiking unit locations is more amenable to downstream tasks such as image registration and visualization (see **Fig. 4, 5**).

### 2.5 Motion inference and registration

Given a set of dense Poisson denoised localization images for each second of recording, we estimate relative motion between frames by treating the set of images as a time-series video and use image registration techniques. Several techniques have been recently introduced to infer motion in Neuropixels recordings. The method proposed in [27] follows a non-rigid template based approach, where an average image of localizations is used as an anchor to register individual localization images using a phase-correlation based displacement estimator [9]. In contrast, the method in [29] follows a decentralized non-rigid approach, treating each localization image as its own template performing decentralized pair-wise motion estimates. This method has been shown to be more robust than the template based approach and we utilize it here to infer displacement both within the NP plane (**z, x**) and orthogonal to this plane (**y**) for the localization images. For other methods that yield only planar localizations, we just estimate vertical (**z**) and horizontal displacement (**y**). The motion estimates for un-denoised and Poisson denoised localization image sets are shown in **Fig. 5D**. Once we estimate motion estimates for each time frame, we then correct for motion by volumetrically (or planarly) translating each image by the negative of the local non-rigid displacement that is estimated for that frame at that location on the probe, bringing all frames into alignment. Example average images before and after alignment can be seen in **Fig. 5H**.

### 2.6 Point-cloud registration

In addition to the image-based registration approach described above, we also experimented with direct point-cloud registration approaches, which skip the formation of an intermediate image representation and are therefore amenable to applications involving higher-dimensional spike featurizations. Representing the spikes in each second of data as a point set, we remove outliers [17] and compress the point cloud via simple hierarchical clustering, parameterized by a single maximum merge distance variable. Then, for each pair of compressed point clouds, we run the iterative closest point (ICP) algorithm [2] to find the displacement in the *z*-direction that minimizes the distance between the clouds, weighted by amplitude and number of spikes. To avoid poor local optima, we utilize a coarse grid search in *z* to initialize the algorithm; see the supplement for full details. The two motion inference methods lead to comparable displacement estimates on the datasets presented in this paper.

## 3 Results

### 3.1 Comparing localizations and clustering

**Fig. 3A**’s first three panels show the {**x, z**} inferred location of detected spikes in 200 seconds of a Neuropixels 2.0 recording, using center of mass (CoM), the Hurwitz et al. [11] method and the new proposed method. (For the Hurwitz et al. method, channels within 35 *μm* from the main channel are included and amplitude jitter is set to 0 *μV*.) While CoM does not allow us to separate waveforms along the **x**-axis or localize outside the convex hull of the electrode, Hurwitz et al. and our method induce spike clusters that appear very different. Hurwitz et al.’s clusters tend to be more isotropic and often appear “paired” on both sides of the probe, while our method leads to differently sized clusters, without pairing. Low amplitude clusters are more spread out than higher amplitude clusters, which is expected as noisier amplitude measurements will lead to noisier localizations.

**Figure 3:**
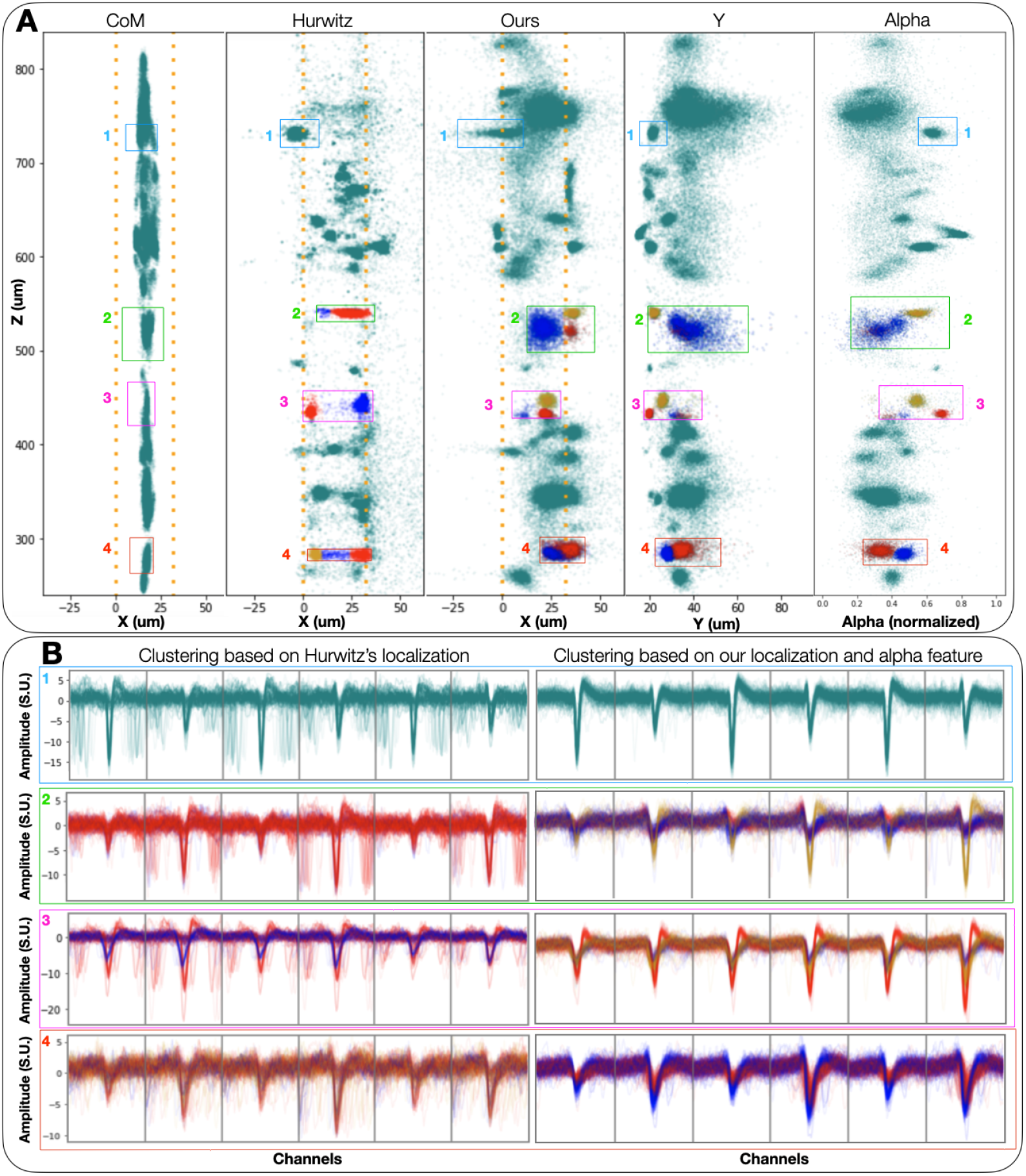
Inferred 3D spatial features yield improvements in waveform clustering. (**A**) {x, z} locations of spikes detected from 200 seconds of NP 2.0 recording, inferred by the center of mass baseline (left), Hurwitz et al.’s [11] state-of-the-art method (second to left), and our method (middle). The two scatter plots on the right of the figure represent z vs. y and *α*, where y and *α* are additional features our method learns. The four colored boxes frame spikes that are shown in panel B, and the yellow, blue and red color of the spikes represent GMM cluster assignments, using {x, z} and {x, z, *α*} as features for Hurwitz method and our method, respectively. The number of clusters has been chosen by hand to reflect the shape of the point clouds in each box. Orange squares represent the position of NP 2.0 recording channels. The boxes in each column are not the same size, as we are trying to match spikes localized in the same z-area, which depends on the localization method. (**B**) Waveforms corresponding to each box and cluster, for both Hurwitz et al.’s (left) and our method (right). (We don’t attempt to cluster based on CoM features, due to the lower resolution of the CoM output.) Note that there is no clear one-to-one correspondence between the left and right waveforms, as our denoising / collision removal can change localization drastically. The many non-centered waveforms in the left column show collided spikes that have not been localized properly as Hurwitz et al. method lacks collision removal. In addition, we find that the Hurwitz et al. approach often spatially separates similar waveforms, while failing to isolate different units. On the other hand, our location-based clusters correspond to visibly apparently separate units, without corruption from poorly localized collided spikes. Moreover, the additional feature *α* can correctly separate different waveforms when {x, z} features alone cannot (red box). See supplementary material for further details.

**Figure 4:**
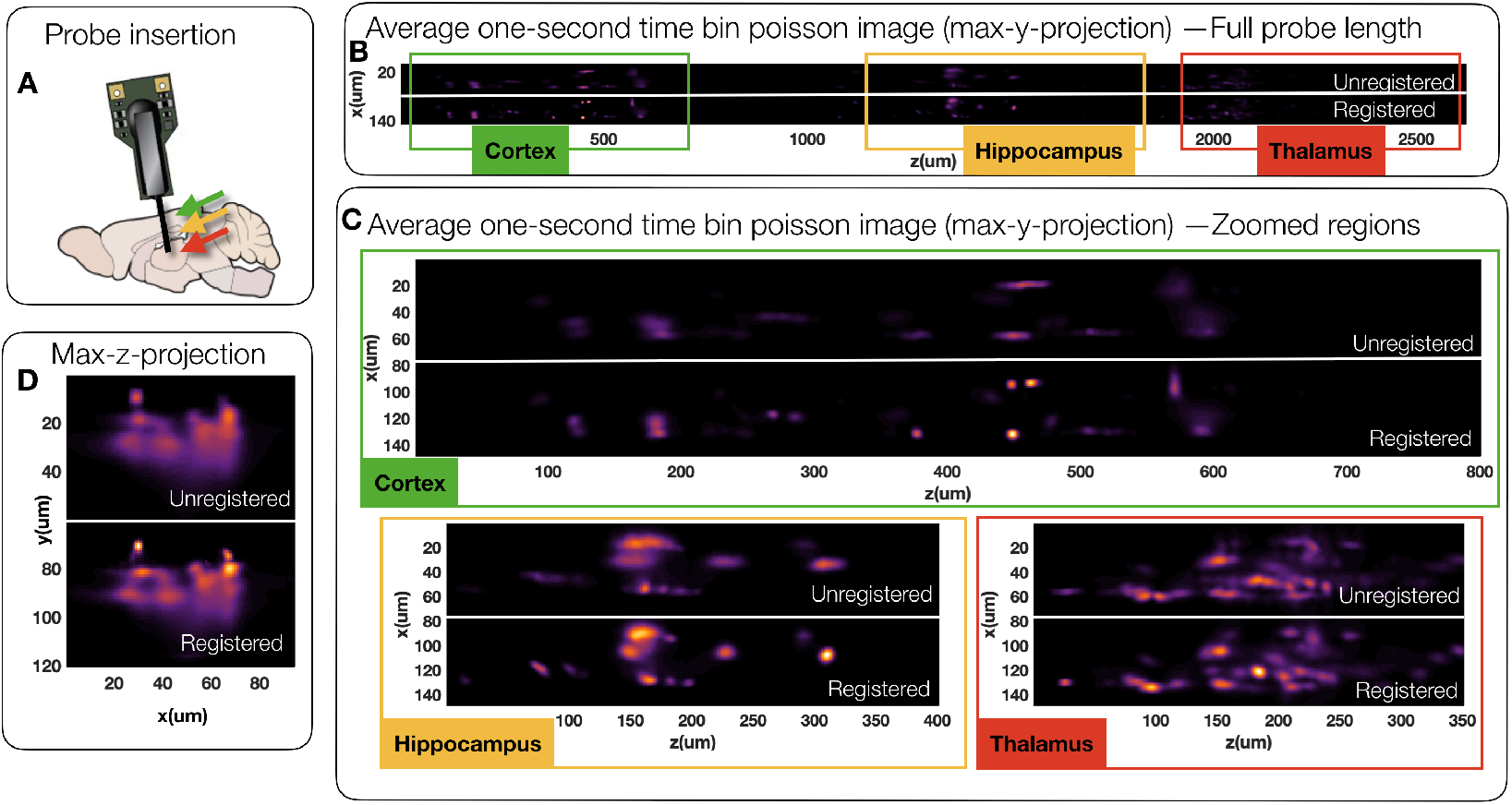
Averaging motion corrected and registered localization images enables a crisp visualization of spiking density. (**A**) We analyze an NP 2.0 recording that has been inserted in the regions of cortex, hippocampus, and thalamus of a mouse brain. The average of thousand one-second localization images after motion correction and registration yields a crisp mapping of neural units in three distinct anatomical regions. Max y-dimension visualization is shown in panels (**B,C**) and max-z-dimension visualization is shown in panel (**D**). (**C**) Zooming into cortical, hippocampal, and thalamic depth regions shows a significant improvement of the crispness of the average localization image after registration. Note that motion effects appear as long blurry streaks in averaged images, whereas well corrected motion yields globular and bright individual clusters. (**D**) Maximum projection along the z-dimension emphasizes localization quality as a function of depth and shows a crisper localization of units near the probe and more uncertain localizations farther away (in depth), owing to higher localization uncertainty for more distant, smaller spikes; this effect is also visible in **Fig. 3A**. See supplementary material and video (video-figure-4.mp4) for a detailed comparison of unregistered/registered frames.

Our method infers two additional features, **y** and *α*. **Fig. 3A**’s two rightmost columns show scatter plots of **z** vs **y** and *α*. From the clusters in boxes 2,3,4 in the right column, it appears that *α* can separate waveform clusters more strongly than **y**, suggesting that this variable contains useful information about the spatial shape of each spike; see the Appendix for further exploration.

The four boxes in blue, green, pink, and red highlight the different spike locations corresponding to similar z-location for each method. For the spikes inside these boxes, we clustered Hurwitz et al. {**x, z**} locations, and our {**x, z**,*α*} features using Gaussian Mixture Model with 2 or 3 components. The corresponding waveforms are shown in **Fig. 3B**. The new method provides visually improved waveform clustering here; the separation induced by *α* is also coherent with the waveform shapes.

A striking difference between the **Fig. 3B** left and right columns’ waveforms is that the right column waveforms appear much less noisy. Most of Hurwitz et al. waveforms appear not centered, but really are collided waveforms representing distinct neural units that are poorly disambiguated. In panels 1, 2, and 3, the Hurwitz et al. method localizes small amplitude spikes collided with large amplitude ones in the same region as the largest amplitude spike. This highlights the importance of using the individual-waveform neural network denoiser. In the supplementary material we contrast the results of all three localization methods using denoised vs raw waveforms. Finally, we find that the Hurwitz method sometimes experiences instabilities in the localization leading to separated clusters in the estimated localization space that do not correspond to separations in the raw data. **Figure 3** panel 4 illustrates this idea, as many spikes of similar amplitude are located by Hurwitz et al. method on either the right or left side of the probe (yellow or red clusters).

### 3.2 Visualizing spike densities after motion correction and registration

After localization and image-based denoising and alignment (as described in the Methods), we compute the average post-aligned image to visualize the volumetric spatial layout of distinct spiking units in the vicinity of the probe. The particular recording that we analyze has been inserted through the cortex, hippocampus, and thalamus of a mouse brain (see **Fig. 4A** for an approximate insertion positioning). The average localization image before and after motion correction and registration for the full probe length is shown in **Fig. 4B**, where we can make out three spatially distinct populations of neurons. Using the depth information and the approximate insertion order, we can then identify and annotate the specific anatomical regions that the neurons reside in and zoom in to observe their densities **Fig. 4B,C**. Note that there is a higher density of spiking units in the thalamus (**Fig. 4C-bottom-right**). By contrasting the unregistered average images with the registered ones, we can see the effects of motion, such as blurred streaks of cell shapes, are mitigated after registration, yielding sharper images of neural populations. Also, visualizing the average image projected along the length of the probe (**z-axis**) shows that while registration does improve sharpness of localization in units close to the probe, “deeper” units that are further away from the probe (and therefore have lower amplitude) remain localized with more uncertainty, as visible also in **Fig. 3A**.

### 3.3 Motion inference and registration evaluation

To further quantify whether our localization and Poisson denoising enable better motion estimation and image registration, we compare the resulting motion estimates and registration metrics using the localization images generated with our method versus the localizations from Hurwitz et al. [11] and the center-of-mass localizer (CoM). To evaluate registration quality, we utilize a Neuropixels 2.0 recording with physically introduced exaggerated motion using an actuator motor (**Fig. 5A,B**). The details of this publicly available recording are shared in [27]. The motion estimates based on all three localization methods and using raw versus denoised localization images can be seen in **Fig. 5D**. The main conclusion is that Poisson denoising significantly improves the motion estimation jitter for all methods compared to utilizing the raw localization images. Furthermore, our localization method affords the estimation of an additional dimension of motion, namely along the depth dimension (**y-axis**) to yield more refined motion estimates. To evaluate motion estimation quality along the length axis (**z-axis**), we provide raster plots before and after registration using all three localization methods after Poisson denoising (**Fig. 5E**). This is essentially a maximum **y-** and **x-** projection of registered images, retaining a singleton dimension **z-** to evaluate residual jitter. The registered rasters using localization images from all three methods are qualitatively well stabilized, with nominal improvements afforded by our localization (highlighted by green arrows in (**Fig. 5E**). Evaluating the registration performance using all dimensions shows that our localization, thanks to its three-dimensional spread of features, provides a better reduction of residual motion (**Fig. 5F,G**) and sharper alignment of images (**Fig. 5H,J,K**) compared to the registration using Hurwitz et al. [11] and CoM localization images.

**Figure 5:**
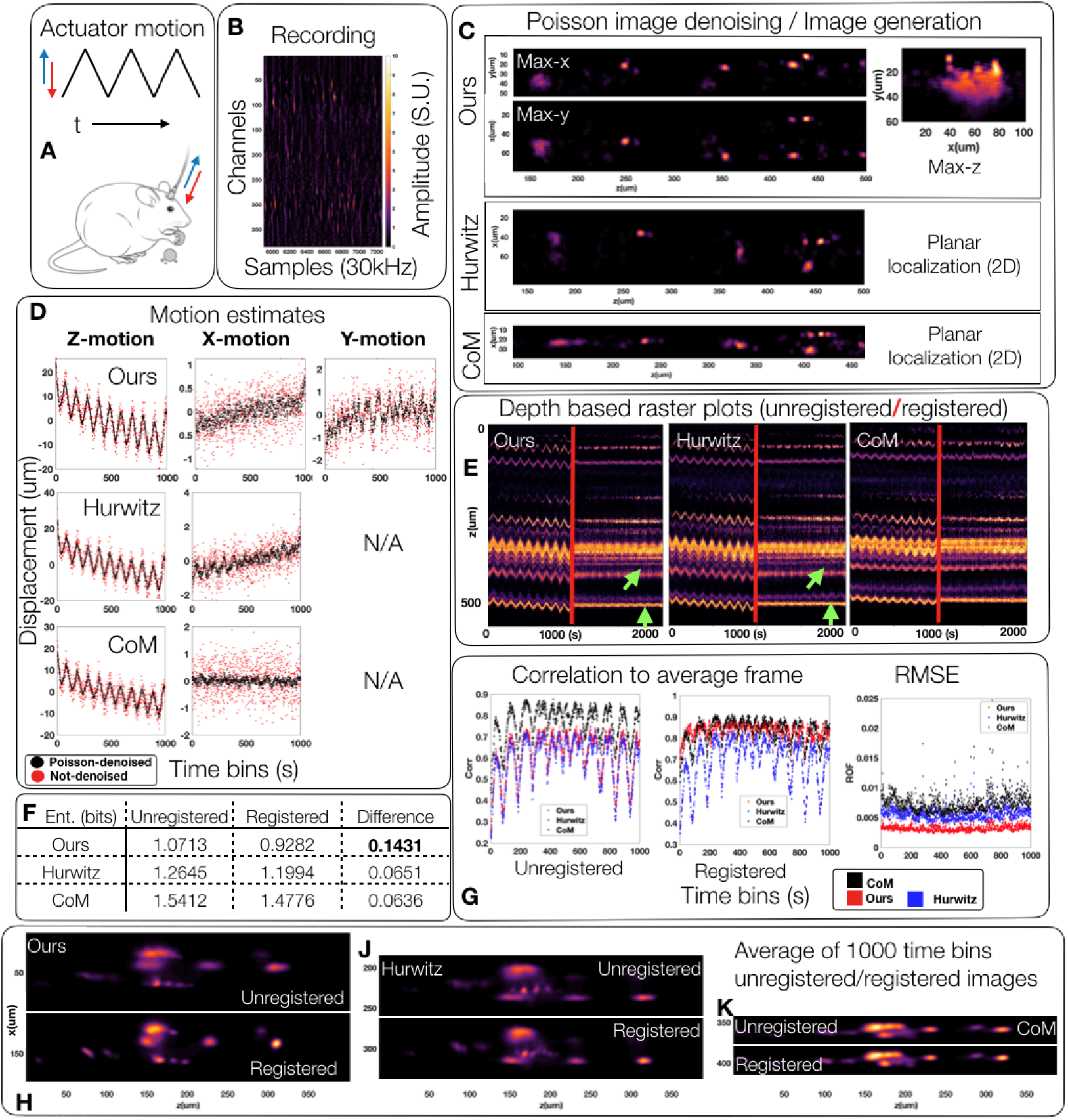
Improved localization enables better motion correction and registration of Neuropixels recordings. (**A, B**) We analyze a mouse NP 2.0 recording that has been subjected to mechanical saw-tooth motion [27]. (**C**) The 3D localization images of spikes after Poisson denoising in a thousand second recording are displayed in comparison to the planar localizations provided by Hurwitz et al [11] and the center of mass technique. Poisson denoising the localization point clouds for all three techniques yields a continuous image representation of spiking unit locations and a one thousand frame video representation (one frame for each second of data). (**D**) We apply the existing registration technique [29] on time-binned image representations of data to estimate the amount of z, x, and y motion for all three localization techniques. We show the motion estimate for each localization technique, with and without Poisson denoising. Poisson denoising significantly improves the noise jitter in motion estimation. (**E**) Visualizing z-direction raster plots of the unregistered and registered recordings (after Poisson denoising) shows stabilization of motion effects for all three methods with nominal improvements by our method over others. Green arrows denote areas of the raster plot that have been well stabilized using our localization versus the localization of Hurwitz et al. (**F, H, J, K**) Visualizing the average image after registration using our localization shows significant decrease in image entropy (as a measure of localization “sharpness”) over compared methods. (**G**) Additionally, our localization affords the highest average correlation of registered images to the average image and the lowest RMSE. Note that CoM method’s high average correlation after registration should be contrasted with its high values prior to registration. Since this localization provides highly blurred images, the average correlation after registration is vacuously high. See supplementary material and video (video-figure-5.mp4) for a more detailed comparisons of unregistered/registered frames using the three localization techniques.

In further detail, we evaluate registration performance using three metrics. First, we compute the average correlation of aligned images to the average image as commonly evaluated in the image registration literature [22]. We contrast this with average correlation before alignment to note the improvement in correlation that alignment provides. Note that the CoM average correlation is high both before and after alignment, denoting that this score is not due to good motion correction but rather to the overall spread of CoM localization that vacuously yields a high correlation to the average image regardless of motion correction. In this metric, our method outperforms both Hurwitz et al. and CoM. We also compute the RMSE of the individual frames to the average frame after registration. Our method also yields the lowest RMSE, showing that individual frames align well to the average frame. Lastly, we evaluate the “sharpness” of the average frame by computing its image entropy. In this regard, our average frame is shown to be qualitatively (**Fig. 5H,J,K**) and quantitatively (**Fig. 5F**) sharper than the results from the compared methods.

## 4 Conclusion

In summary, we provide a simple denoising and point-model triangulation approach to infer three-dimensional source locations from individual spikes in Neuropixels recordings [13, 27]. This localization is shown to facilitate better clustering of distinct neural units as well as enabling better estimation and correction of motion in datasets where the spiking units experience non-stationarities due to probe drift. We also demonstrate the benefits of converting the spiking unit localization point sets into continuous images using Poissonian denoising [18], enabling informative visualizations of the neural source distributions in the vicinity of the Neuropixels probe.

Two open directions are clear for future work. First, as emphasized above, we have used a highly simplified point model to perform localization here. Elaborations of this model should lead to improved accuracy, though robustness and speed of the resulting localizer are also critical for downstream applications and should be balanced accordingly. Second, establishing the accuracy of the proposed methods experimentally will be a challenging but critical next step.

## Broader impact

Neuropixels probes are a recently-developed neurotechnology that enable us to record from hundreds of neurons simultaneously in multiple regions of the brain. Our work will improve scientific conclusions derived from Neuropixels recordings; downstream, we expect related methods to also improve the performance of brain machine interfaces based on extracellular neural recordings.

## Acknowledgments

We thank Nick Steinmetz for collecting and sharing Neuropixels recordings, Olivier Winter and the International Brain Lab for testing our methods, Cole Hurwitz and Matthias Henning for helping us draw comparisons with their localization method, as well as Alessio Buccino and Samuel Garcia for helpful discussions and references. We also thank Cyrille Rossant for providing support for the Datoviz platform. This work was supported by the following grants: Gatsby Charitable Foundation GAT3708, NSF DBI-1707398, NSF 1546296, NIH U19NS104649, NIH U19NS123716, Simons Foundation 543023, Wellcome Trust 209558, and Wellcome Trust 216324.

## Supplementary material main contributions

1. Details on the localization, motion estimation and registration procedures, as well as a discussion on the importance of our denoising step
2. An illustration and evaluation of the localization method on a synthetic toy dataset
3. Comparison between our 3d localization features and waveforms’ shape features
4. Neuropixels 1.0 localization and registration results
5. Videos to illustrate the performance of our improved registration method

## 1 Localization and optimization

### 1.1 Localization and optimization details

The individual waveform neural network denoiser we use to get clean waveforms and unbiased amplitudes is defined in YASS [8]. For each type of probe, the neural network can be trained following instructions at https://github.com/paninski-lab/yass/wiki/Neural-Networks---Loading-and-Retraining.

This neural network is an individual waveform denoiser. It denoises the signal on each channel separately. When trained, it expects the input waveform’s minimum to be found at a timepoint close to a given value. For example, if trained with waveforms that have their minima around timepoint 41, it will return a clean waveform that takes its minimum close to timepoint 41 as well. It uses this information to remove collisions: If a waveform takes its minimum around 60, it will be automatically treated as a collision and the output will be a waveform taking its minimum around 41. For good accuracy across channels, we upsample and align waveforms before denoising to correct for micro-time shifts between channels. This process removes “close” collisions. However, if a “far away” neuron (for example, localized at a very different z-position) fires simultaneously, the signal will be equal to the sum of the collided waveform and noise, and the denoiser will return the denoised waveform instead of detecting it as a collision. It is important to discard this waveform when computing localization.

To remove these “far away” collisions, we need to first run a de-duplication step (implemented in many spike sorters such as YASS [8] and Kilosort [10]), to get an estimate of the localization and its main channel, giving a set *C*_*m*_ of relevant channels.

For each denoised waveform *w*_*n*_, we find its max channel 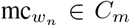 and perform localization using the denoised amplitudes recorded at channels 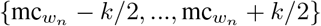 by minimizing 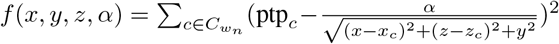 over {*x, y, z, α*}. We optimize this function using least-squares optimization after using center-of-mass as a simple quick initialization. The function we optimize is non-convex, but we show in section 1.2. that this optimization method is suited to the task. The whole procedure is summarized in Algorithm 1.

**Algorithm 1.**
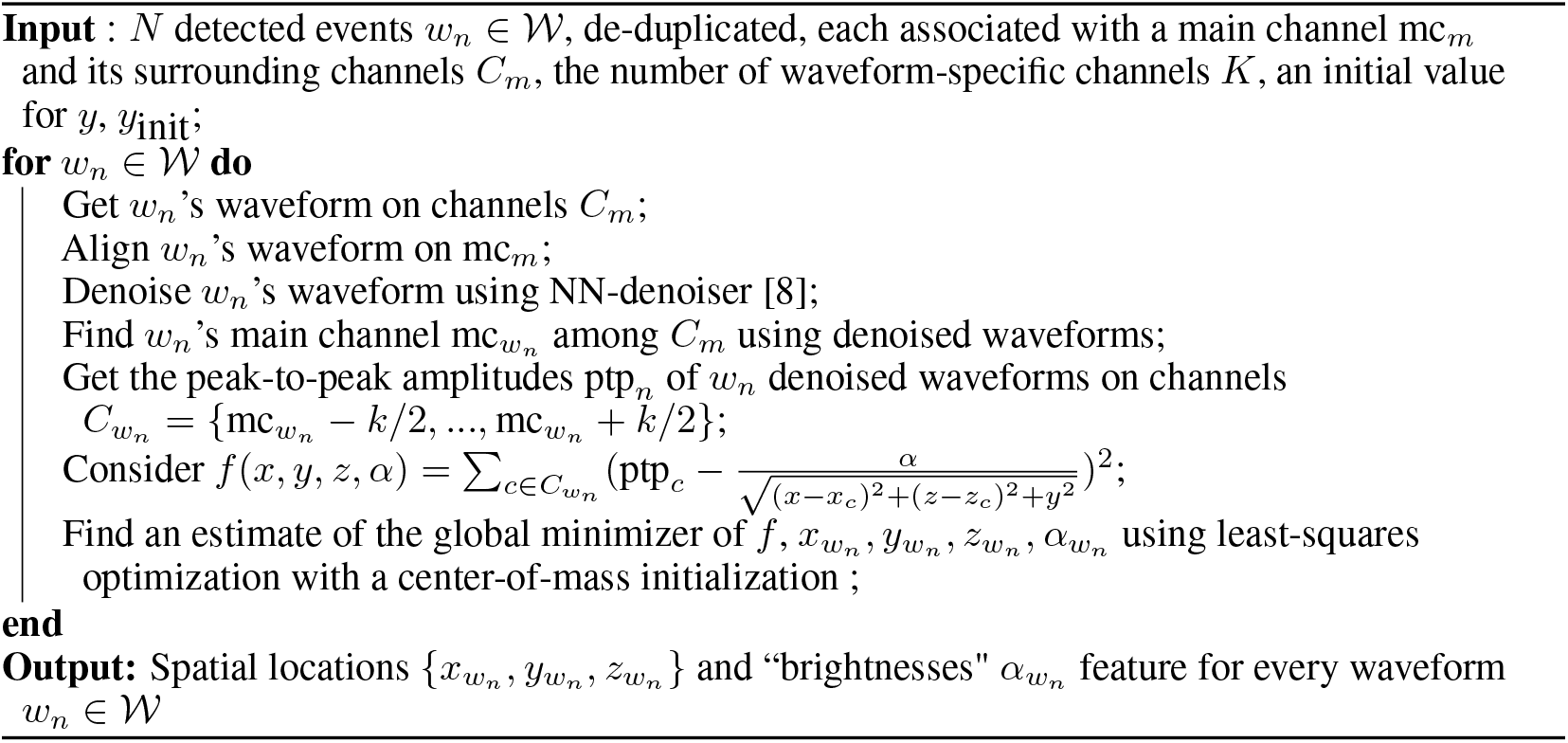
Localization.

*k*, the number of channels used for localizing denoised waveforms, is the only hyperparameter of our model. Choosing *k* to be small will lead to “flat” clusters (exhibiting small standard deviation along z-axis and high standard deviation along x-axis), and loss of precision along the z-axis, while choosing a large *k* will allow small amplitude, noisy channels to be taken into account when localizing, which will spread out the clusters and lead to a loss in precision. The set of “relevant” channels *C*_*m*_ has to be larger than *k*.

### 1.2 Evaluating optimization on a toy dataset

We validated the method on a toy dataset. We reproduced the position of 80 channels, for both NP1.0 and NP2.0, and randomly assigned {*x, y, z*} positions of 1000 neurons drawn from a Gaussian distribution. We then fix *α* = 500 and compute the corresponding amplitudes on each channel using 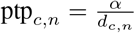 where *d*_*c,n*_ is the distance of {*x*_*n*_, *y*_*n*_, *z*_*n*_} to channel *c*. We use our localization method to infer the corresponding locations, and show that our model can accurately recover {*x*_*n*_, *y*_*n*_, *z*_*n*_} locations (**Fig. 1**).

### 1.3 Reliable estimation of spikes amplitudes using the neural network denoiser

To explore the quality of our amplitude estimation using the neural network denoiser described in section **2.2**, we selected 170 clean templates obtained after running YASS [6] on a Neuropixels 2.0 dataset, added background signal taken randomly from the same dataset to get simulated waveforms, and computed amplitudes before and after denoising. **Fig. 2** shows a scatter plot of the true template amplitudes vs. the waveform amplitudes (in blue) and the denoised waveform amplitudes (in red). Denoised amplitudes are much closer to the true amplitudes of the templates, as desired.

### 1.4 Localizations without denoising

To further illustrate the importance of the denoising step on localization, **Fig. 3** (analogous to **Fig. 3** in the main text) shows the inferred features and associated GMM clusters for localizations without denoising. The clusters of locations appear noisier and more spread out. Moreover, we see that many collided spikes belonging to different units are located in single clusters, since there is no way to disambiguate collisions with the amplitude-based localization method.

**Figure 1:**
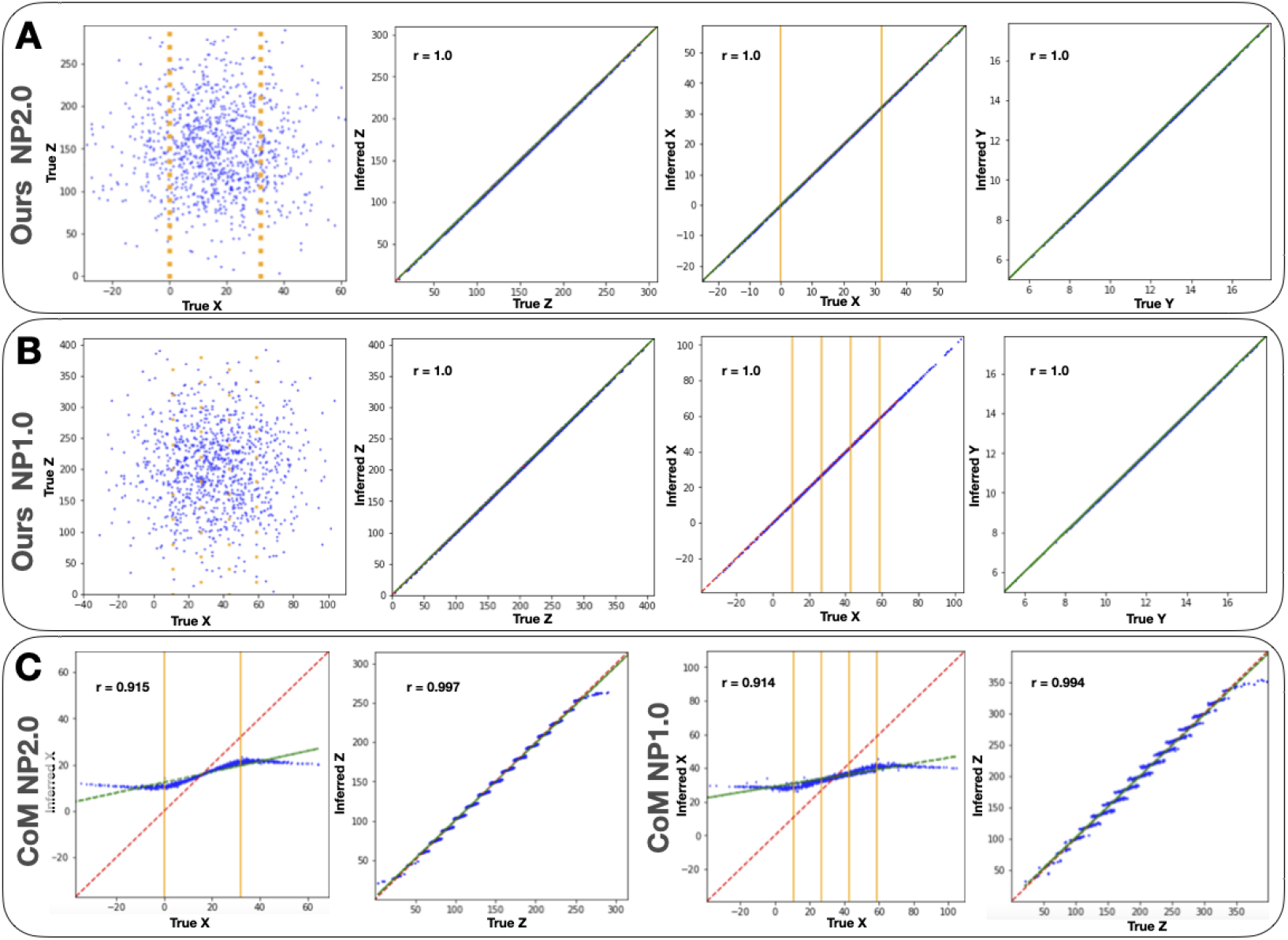
Simulated toy data localization. We sampled *x, z, y* spike locations from a Gaussian distribution, and computed the amplitudes on each channel using the simple point model. We compare the true locations vs. our model’s inferred locations. (**A**) shows results with channels following Neuropixels 2.0 geometry. Left shows the simulated {*x, z*} locations (blue) and the channel positions (orange). The three scatter plots show inferred {*z, x, y*} vs true {*z, x, y*}. Dashed red line corresponds to “*x* = *y*” and the green one to the regression line. Correlation coefficients are reported on the top left of each scatter plot. The orange lines on the third scatter plot indicate the *x* position of the channels. Panel (**B**) is similar with channels reproducing the geometry of Neuropixels 1.0 probes. Panel (**C**) shows similar scatter plots for the inferred {*x, z*} positions of Center of Mass method, for both Neuropixels 2.0 (left) and Neuropixels 1.0 (right). Our method recovers {*x, y, z*} positions accurately. The Center of Mass method fails at recovering {*x, z*}; it localizes inside the convex-hull of the electrode locations, and “shrinks" positions to the center of the probe. The stair-like pattern on the *z* scatterplots indicate that it tends to localize very close to the channels. We did not compare with Hurwitz et al. method [5] as it does not rely on amplitudes only for localization.

**Figure 2:**
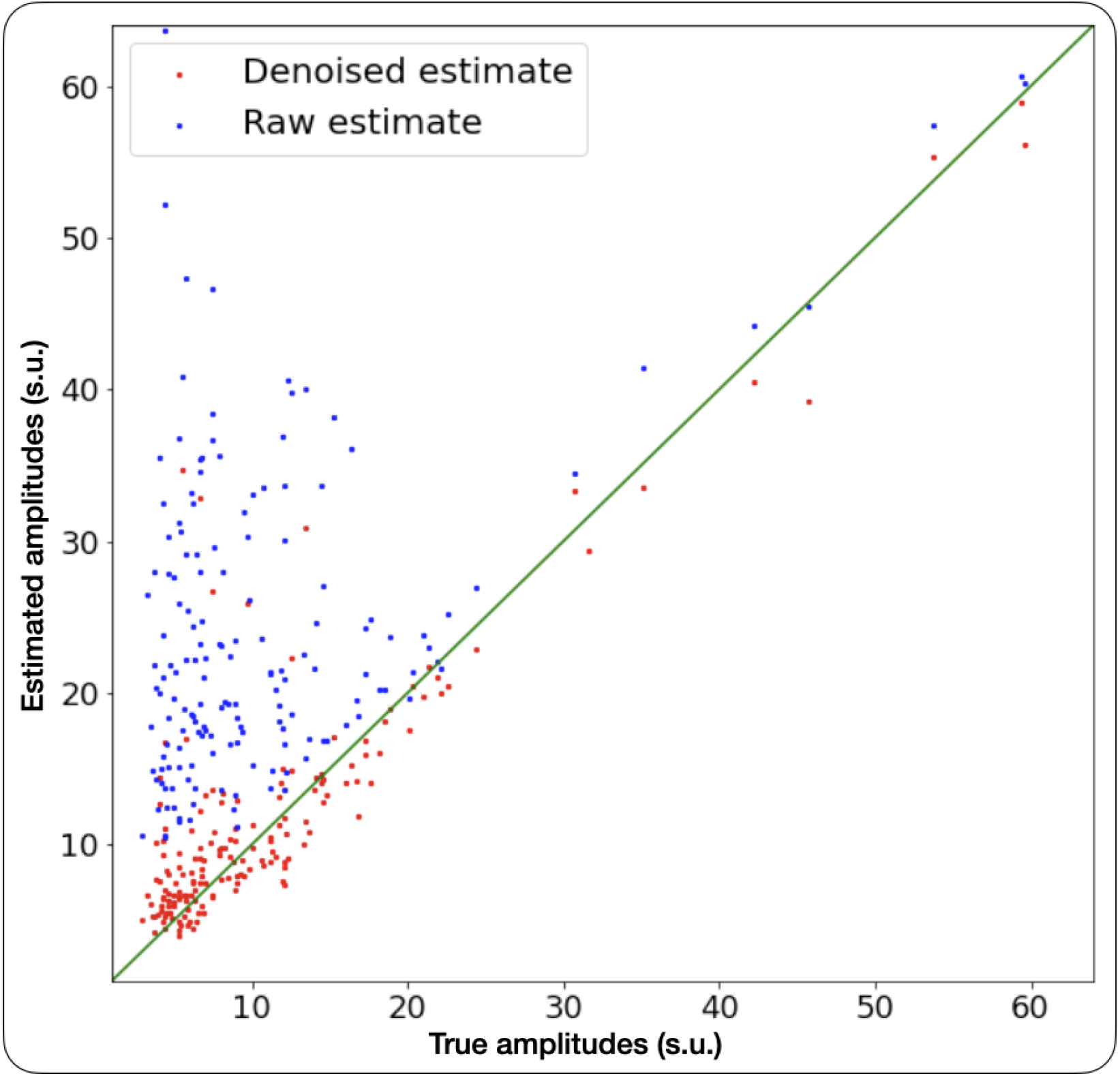
Denoiser’s output waveforms have close to true amplitudes. Scatter plot of the true template amplitudes vs simulated waveform amplitudes (in blue) and the denoised waveforms amplitudes (in red). (See text for simulation details.) Green line indicates *y* = *x*. Overall, denoised amplitudes are much closer to the true amplitudes.

**Figure 3:**
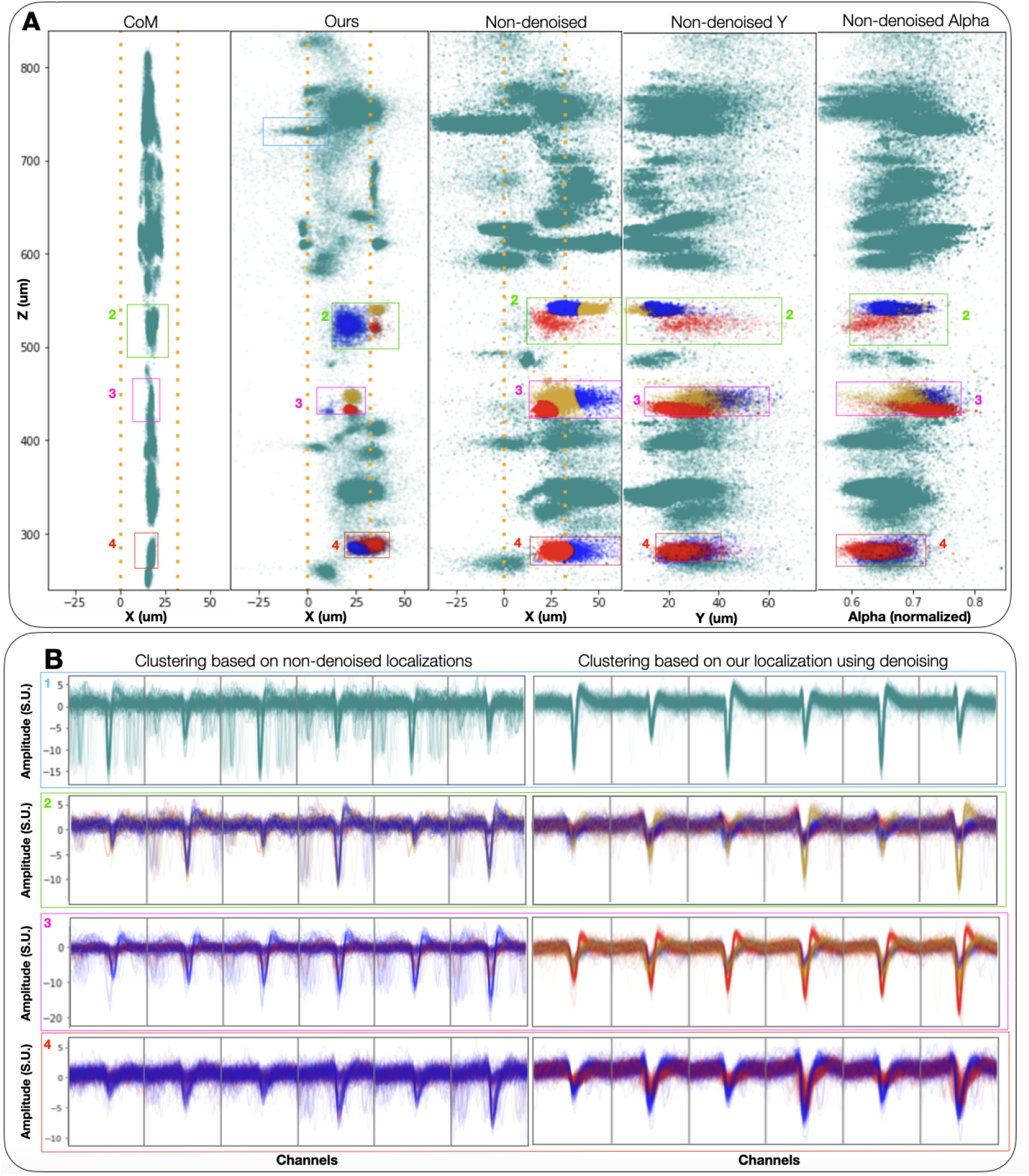
Denoising lead to sharper clusters and disambiguate collided spikes. We use the same convention as the main text Fig. 3 in this figure. (**A**) {x, z} locations of spikes detected from 200 seconds of NP 2.0 recording, inferred by the center of mass baseline (left), our method (second to left), and our method without the denoising step (middle). The two scatter plots on the right of the figure represent z vs. y and *α*. The four colored boxes frame spikes that are shown in panel B, and the yellow, blue, and red color of the spikes represent GMM cluster assignments, using {x, z, *α*} as features for our method with and without denoising. The number of clusters has been chosen by hand to reflect the shape of the point clouds in each box. The boxes in each column are not the same size, as we are trying to match spikes localized in the same z-area, which depends on the localization method. (**B**) Waveforms corresponding to each box and cluster, for our method with (right) and without (left) denoising. The many non-centered waveforms in the left column show collided spikes that have not been localized properly without denoising.

## 2 Comparing features

Figure 4 shows a comparison of our localization features {*x, y, z, α*} to the first two principal components of the waveforms, and an additional “spread” feature, equal to the trace of the covariance of the distribution of each spike amplitudes across channels. This feature should be informative of *y* location, as the maximum amplitude can’t help distinguish between a low-amplitude, close spike and a high-amplitude, far away spike.

Some clusters are better separated by *α* than any other features, indicating that it contains information about the shape or location of the waveforms that is not contained by other single features.

**Figure 4:**
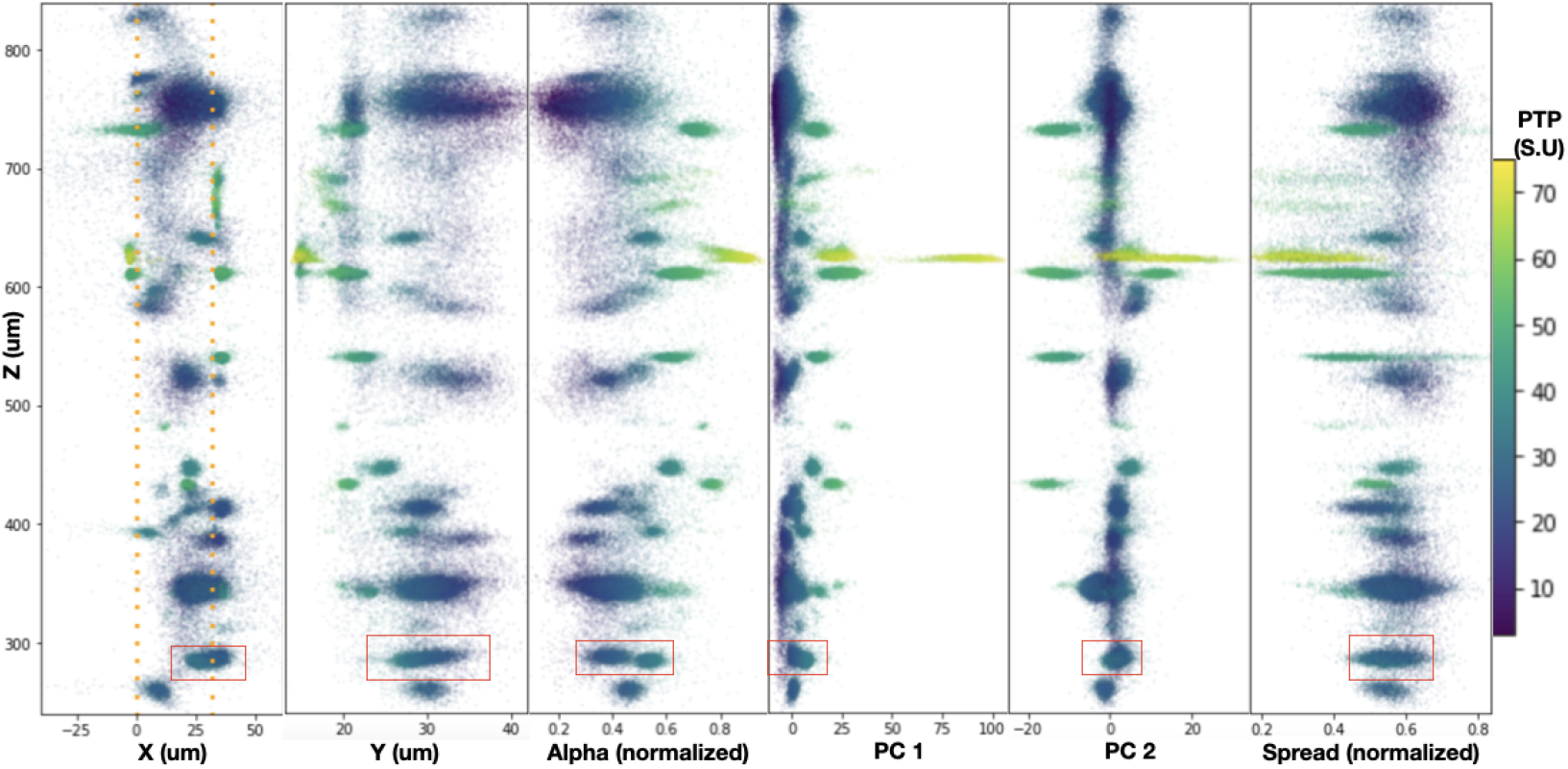
Comparing localization and shape features. Scatter plots showing our inferred {*x, z*} locations of spikes detected from 200 seconds of NP 2.0 recording (left), {*y, z*} locations (second to left), {*α, z*} (third to left), the first two Principal Components of the waveforms vs. *z* (fourth and fifth to left), and the spread of each waveform vs. *z* (right). Spikes are colored by maximum peak-to-peak amplitude. The spread of each waveform is calculated as the covariance of the distribution of the denoised amplitude over the set of selected channels. We expect spread to be informative about *y*, as a far away high-amplitude spike will be seen on multiple channels with low detected amplitude, giving high spread, whereas a small or high amplitude spike close to the channels will correspond to a low spread value. *Y* is determined by the amplitudes. The point clouds inside the red box (around depth 300) are only separated by *α*, and not by the Principal Components of the waveforms, the spread or the maximum amplitude, suggesting that *α* contains additional information useful for clustering. This figure shows scatter plots corresponding to the same spikes as the main text Figure 3, and the red box corresponds to Box 4.

## 3 Comparison of image denoising techniques

Due to the limitations of the point neuron model and the sparse number of observations, the localization methods provide a partial picture of the spatial layout of spiking units. To generate a dense localization images for image based registration, we utilize three different denoising techniques: 1) Gaussian smoothing, Poisson denoising [9] (ours), and Deep interpolation (DI) [7].

- Gaussian smoothing involves blurring the localization images by a Gaussian filter to infill gaps. Here we used a kernel size of 5*μ* m.
- DI is a neural-network based denoising algorithm that takes noisy samples from the original raw data as inputs to train a spatio-temporal nonlinear interpolation model. We applied deep interpolation to a small patch of NP 2.0 raster image (depth 70-582*μ*m and time 1-512s), with one of the network architectures provided by the authors (“unet_single_256”). The network is trained with 5 steps per epoch (total of 7 epochs), with batch size of 1 and “pre_post_frame” set to 1.
- Poisson denoising models the observed localization image as having been corrupted by Poisson salt-and-pepper noise, with the likelihood of observing a spike proportional to its amplitude. Since this process models the noise variance to be proportional to its mean, we first apply the Anscombe transformation [1] to stabilize noise variance prior to denoising the transformed localization image by BM3D [4] or non-local-means [3]. After the transformed image is “denoised” we apply the inverse Anscombe transformation to yield the “Poisson denoised” localization image.

The three approaches to image denoising have slightly different effects on downstream motion estimation. Figure 5 shows the comparison of the probe displacements for the first 512 seconds, estimated by localization images denoised by Gaussian filtering, DI, and Poisson denoising. The estimates are similar, but there is a drastic difference in the run-time (178 for Deep Interpolation vs. 0.27 seconds for Poisson denoising), suggesting that our Poisson denoiser is both effective and efficient. Furthermore, the motion estimate curves yielded by DI outputs of localization images shows dampened peak-to-trough amplitudes of motion in the simulated NP 2.0 datasets, indicating that DI may be slightly over-smoothing the images, obscuring fine details that may be useful for precise motion estimation.

**Figure 5:**
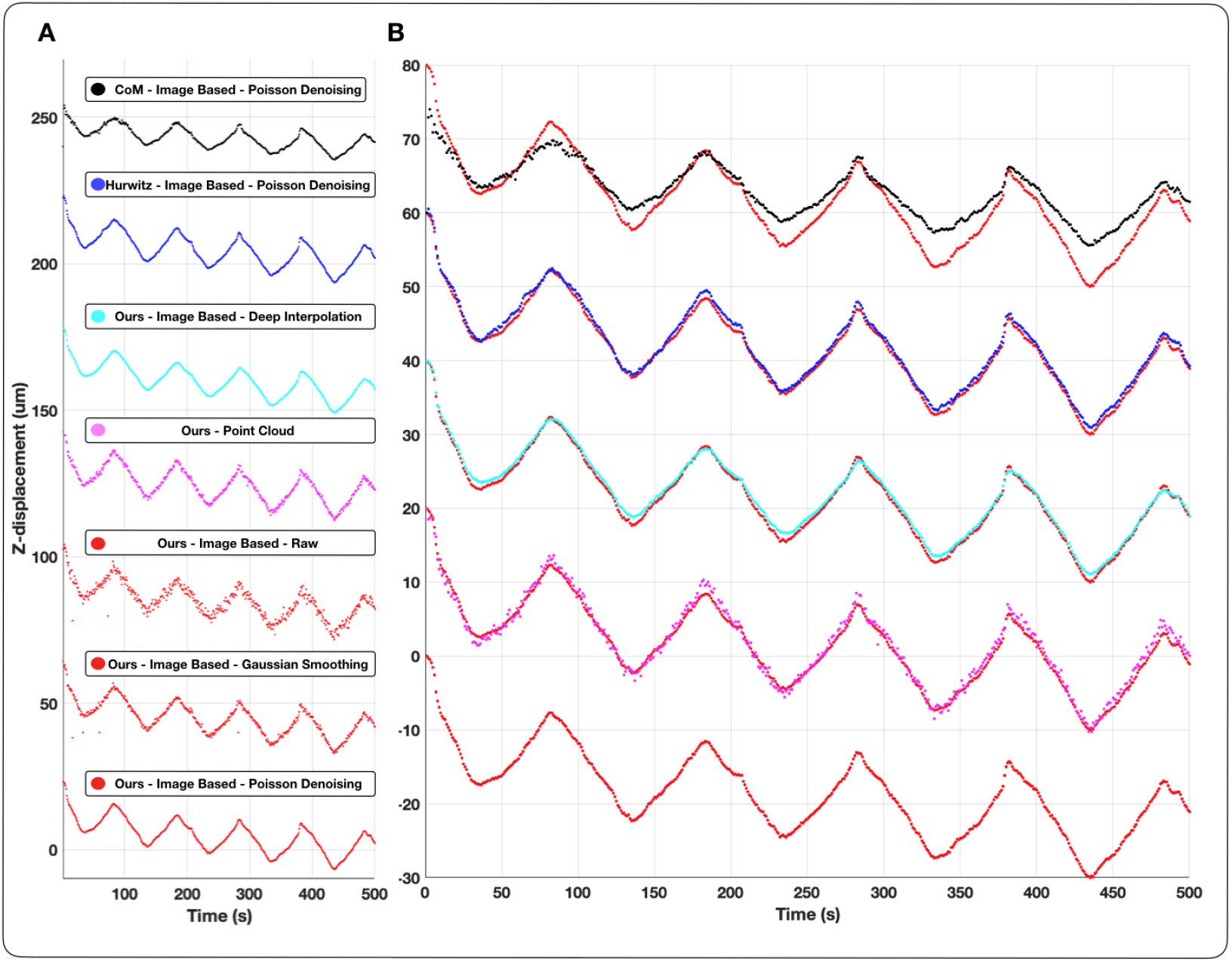
Comparing the effects of different image denoising techniques for motion estimation. **A:** The displacement estimates obtained from our localization images without denoising as well as Poisson denoising, Gaussian smoothing, and Deep Interpolation denoising, compared with the displacement estimates obtained using Center of Mass localization and Hurwitz method of localization with Poisson denoising. **B:** The displacement estimates are superimposed on top of the estimates from our localization method + Poisson denoising to examine subtle differences between the methods. Displacement estimates using our localization method yield similar outputs whether we use image based registration or point cloud based registration. Our localization with Poisson denoising yields slightly “peakier” displacement estimates versus denoising with Deep Interpolation. Gaussian blurring yields noisier estimates of displacement than the other denoising methods.

## 4 Additional details on point-cloud registration

Algorithms 2 and 3 outline the detailed steps of the point-cloud registration technique (in the rigid case, for simplicity), with Figure 6 providing an illustration.

**Algorithm 2.**
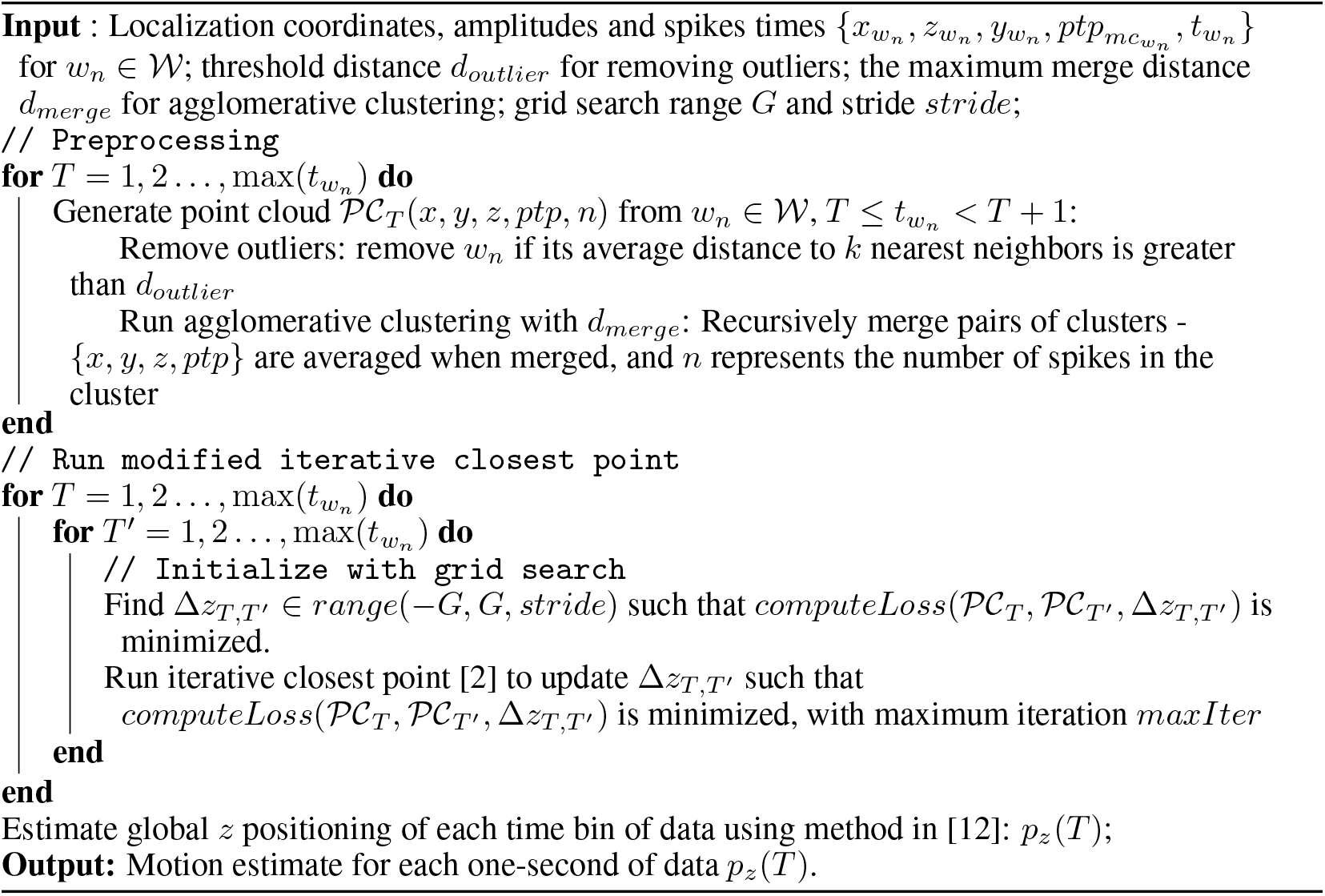
Point-cloud registration.

**Algorithm 3.**
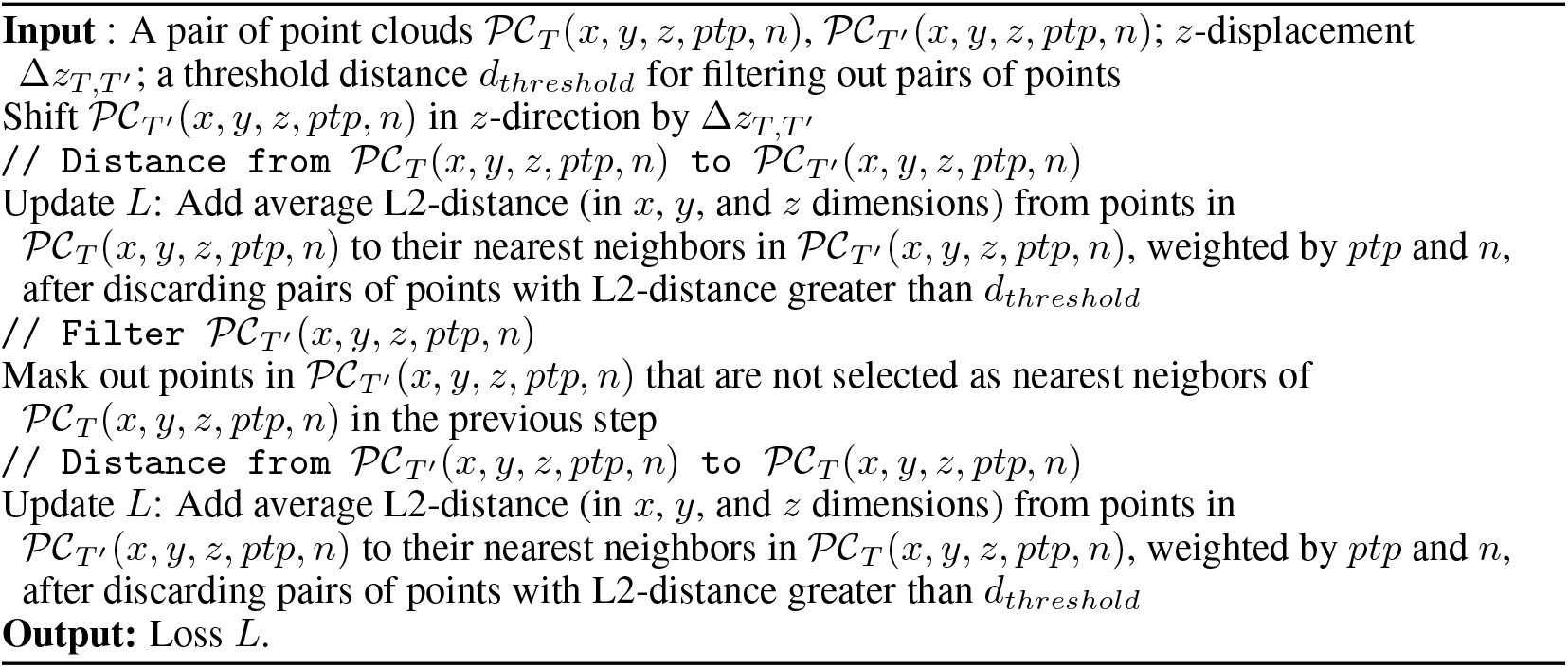
computeLoss.

**Figure 6:**
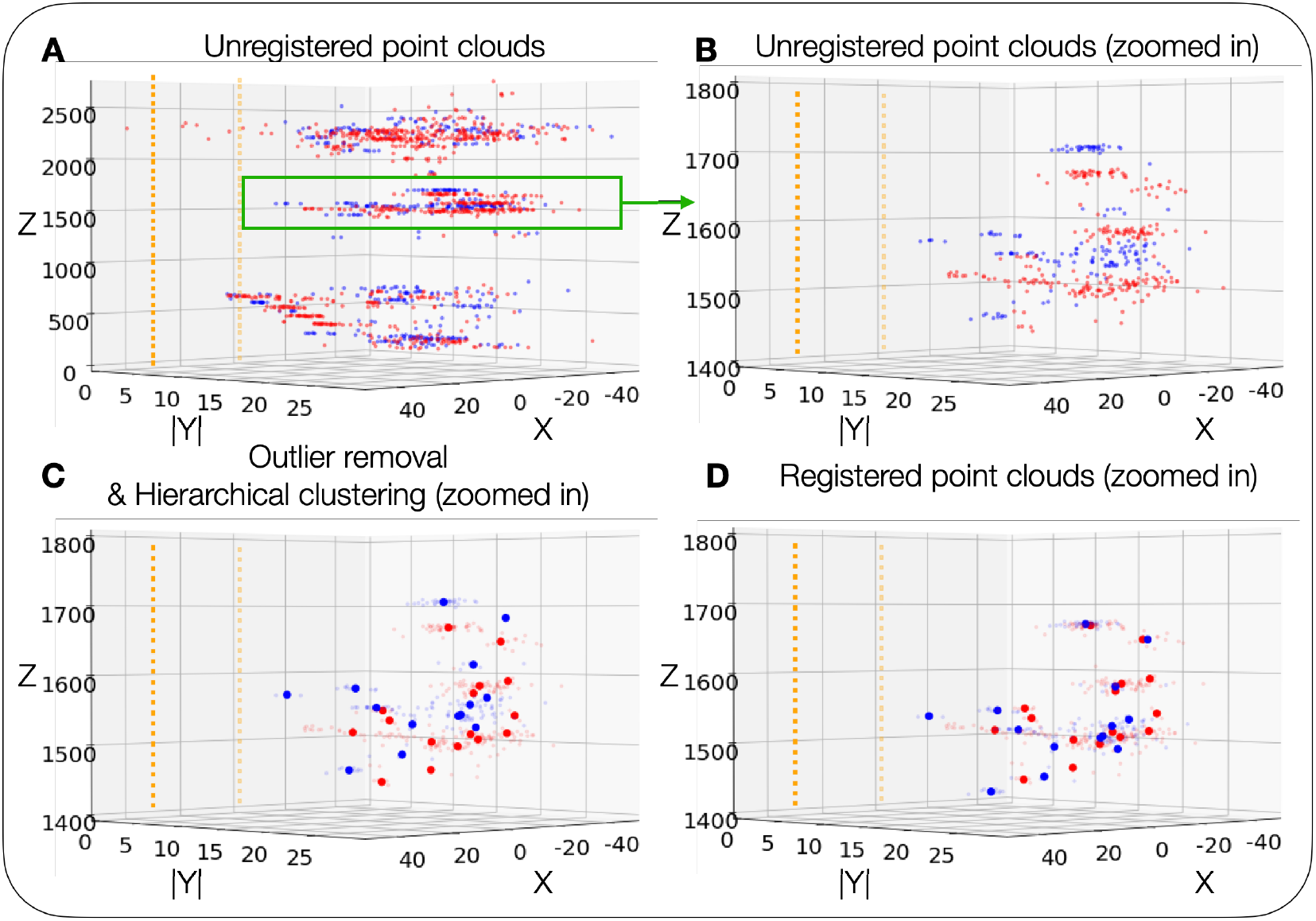
Visualization of point-cloud registration on NP 2.0 recording. Visualization of unregistered point clouds (**A**), unregistered point clouds zoomed in to depth 1400 - 1800*μ*m (**B**), compressed (i.e. outlier removal & hierarchical clustering) point clouds (**C**), and registered compressed point clouds (**D**). Colors represent which second of data each spike is from (red: 1st second, blue: 850th second). Orange squares represent the recording sites of the NP 2.0 probe. Smaller dots represent the original spikes, and larger ones represent the means of the hierarchical clusters. Here we register the blue point cloud to the red one by shifting 33.04*μ*m in the *z*-direction, and we see that the two point clouds overlap to a greater extent after registration.

## 5 Neuropixels 1.0 results

### 5.1 Spike localization

To demonstrate that the localization model is robust to different types of probes, we show improvements over previous localization methods on a Neuropixels 1.0 dataset. **Fig. 7** shows locations inferred by Center of Mass, Hurwitz et al. [5], and our method, and corresponding clusters obtained using Gaussian Mixture Model on the location features. For the Hurwitz et al. method, channels within 50 *μm* from the main channel are included (10 observed amplitudes) and amplitude jitter is set to 0 *μV*.

### 5.2 Motion estimation and registration

We evaluate registration performance to demonstrate that motion estimation is robust to the localizations that arise from Neuropixels 1.0 probes geometry. The quantitative and qualitative results are shown in **Fig. 8**.

**Figure 7:**
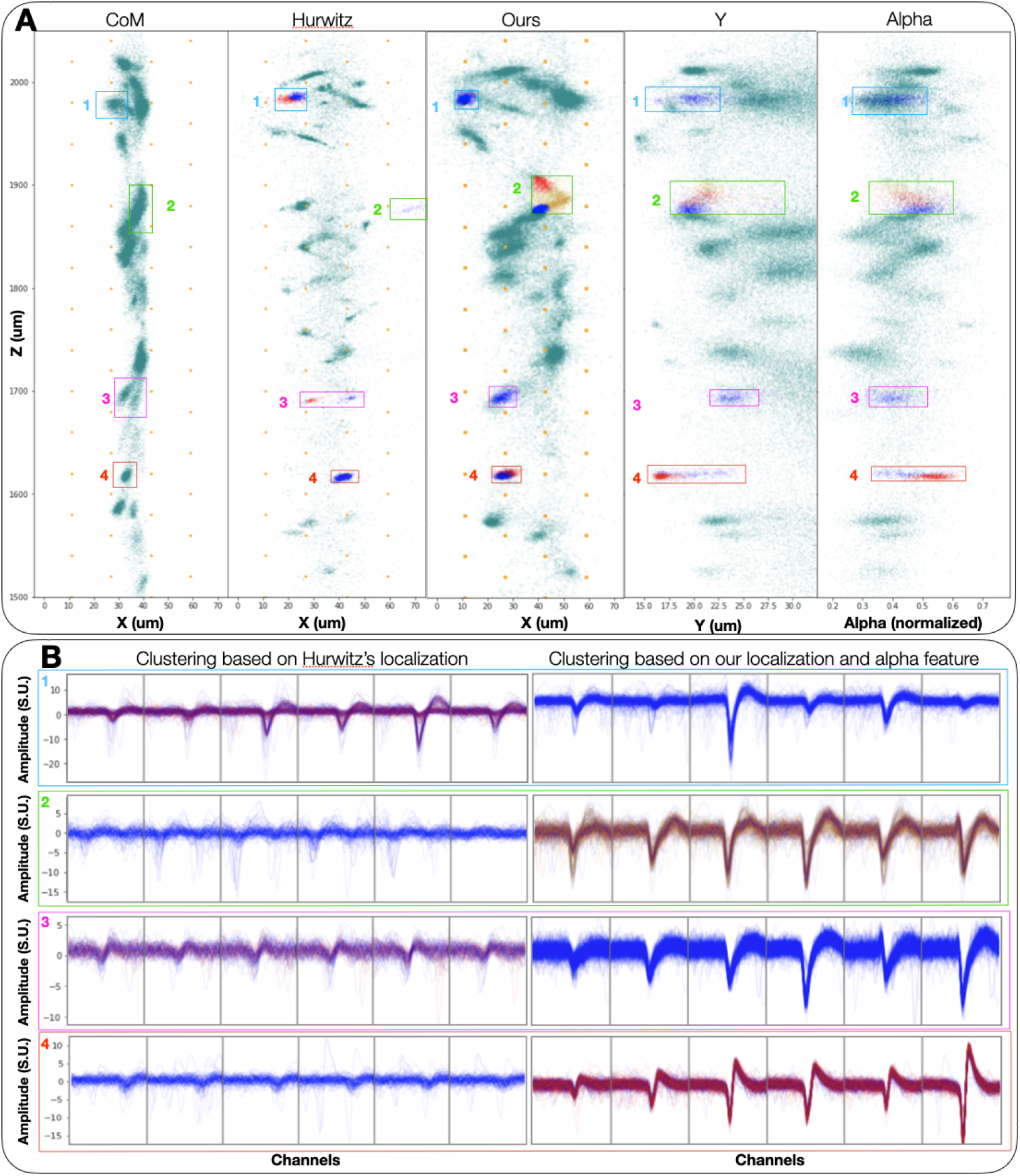
Inferred 3D spatial features yield improvements in Neuropixels 1.0 waveform clustering (analogous to figure 3 in main text) (**A**) shows center of mass, Hurwitz et al. and our localization results, with added features *Y, α* for our method. Spikes in each box correspond to the waveforms in panel (**B**), with blue, red and yellow colors corresponding to Gaussian Mixture Model clustering with the number of components reflecting the cloud points shape. The many non-centered waveforms in the left column of Panel (B) show collided spikes that have not been localized properly by the Hurwitz et al. method, which often spatially separates similar waveforms, while failing to isolate different units. On the other hand, our location-based clusters correspond to similar units for boxes 1 and 3, without corruption from poorly localized collided spikes. Nonetheless, some units here show signs of oversplitting (e.g., boxes 2 and 4), indicating room for potential further improvement.

**Figure 8:**
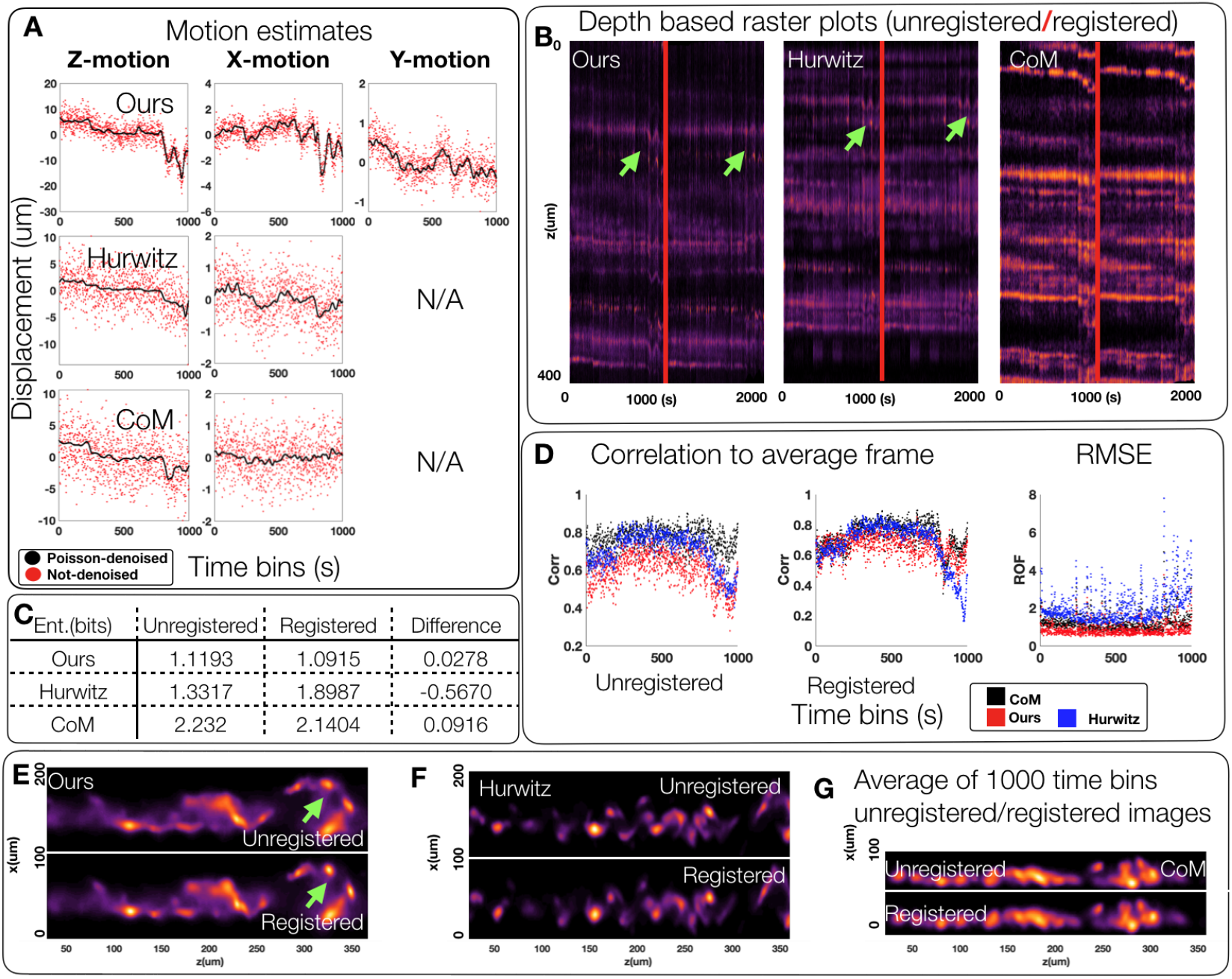
Improved localization enables better motion correction and registration of Neuropixels 1.0 data (analogous to figure 5 in main text) (**A**) We apply the existing registration technique [12] on time-binned image representations of data to estimate the amount of z, x, and y motion for all three localization techniques. We show the motion estimate for each localization technique, with and without Poisson denoising. Poisson denoising significantly improves the noise jitter in motion estimation. (**B**) Visualizing z-direction raster plots of the unregistered and registered recordings (after Poisson denoising) shows stabilization of motion effects for all three methods with nominal improvements by our method over others. Green arrows denote areas of the raster plot that have been well stabilized using our localization versus the localization of Hurwitz et al. (**C, E, F, G**) Visualizing the average image after registration using our localization shows significant decrease in image entropy (as a measure of localization “sharpness”) over compared methods. (**D**) Additionally, our localization affords the highest average correlation of registered images to the average image and the lowest RMSE. Note that CoM method’s high average correlation after registration should be contrasted with its high values prior to registration. Since this localization provides highly blurred images, the average correlation after registration is vacuously high.

## 6 Supplementary videos

### 6.1 Supplementary Datoviz video for figure 1 in main text (Web link: video-figure-1.mp4)

Datoviz [11] is a high-performance interactive data visualization library that we use to visualize our localizations and spike sorting output along the probe, and inspect the clusters in 3-d. In this example video, we are showing four different panels. The first one displays the spikes’ 3-d locations {**x, y,z**} along the probe, colored by post spike-sorting clusters. The second panel shows the same features, colored by maximum amplitude. The third one shows the same features colored by *α* feature, while the last one displays {**x, z**,*α*} along the probe, colored by maximum amplitude.

This interactive plot allows us to scroll along the probe, zoom to inspect clusters (do the spike-sorting units have well-defined clusters, do they correspond to two different clusters, is one cluster separated in two different units?), aggregate spikes in time, and look at different time points in the recordings. The latter provides a good way to visualize the 3-d displacement of the probe.

### 6.2 Supplementary video for figure 4 in the main text (Web link: video-figure-4.mp4)

This video shows the zoomed regions of the Neuropixels probe that correspond to the anatomical regions of cortex, hippocampus and thalamus. Screenshot of this video is shown in figure 9. In each frame top left panel shows the sparse localization image of detected spikes and their spatial positions after having been projected along the depth (y) axis. Top right panel is analogous to top left panel but shows the projection along the horizontal axis (x). Middle panels show the localization images after having gone through Poisson denoising [9]. Bottom panels show the Poisson denoised images after motion estimation and registration.

**Figure 9:**
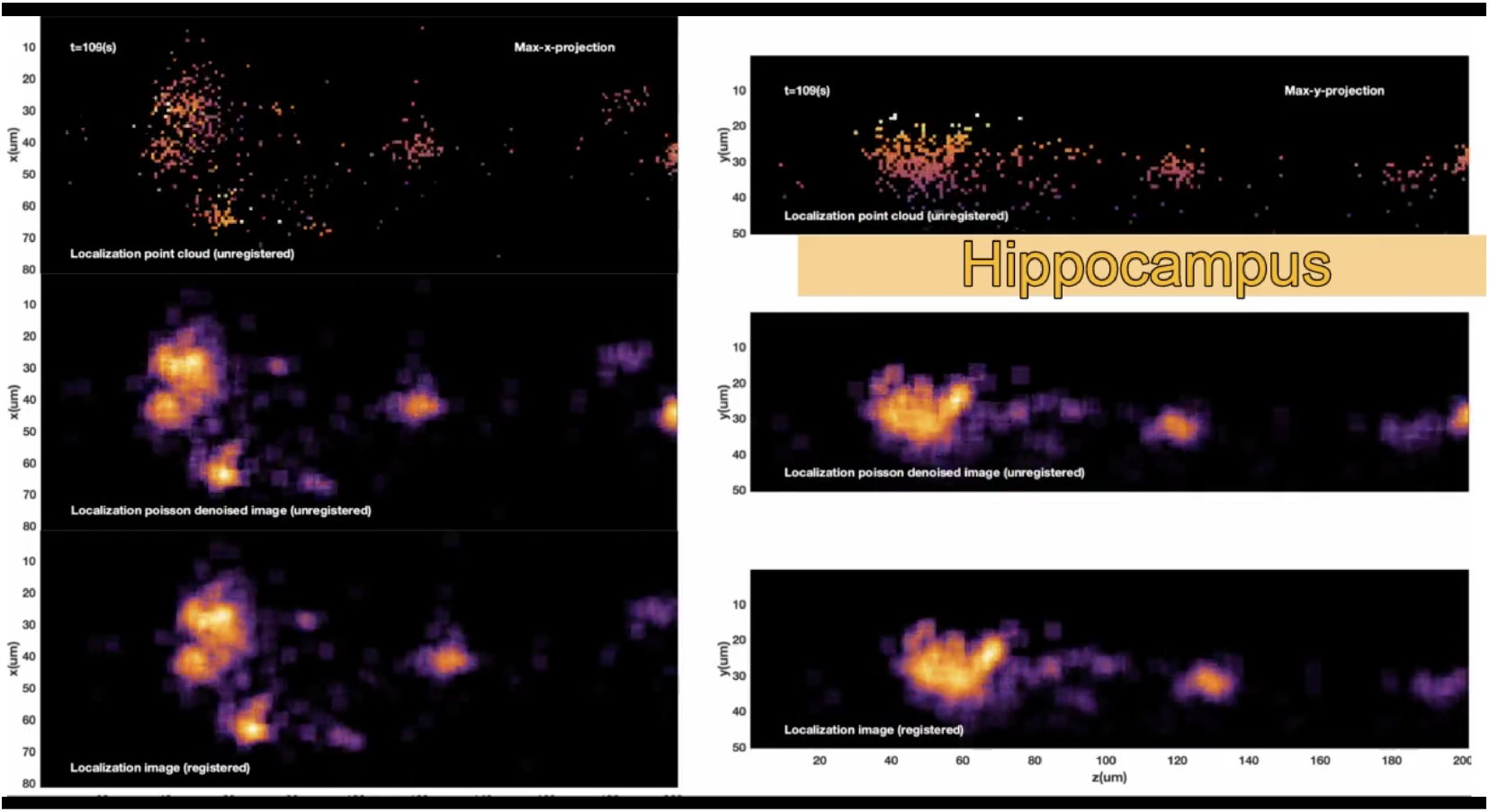
Video representation of Poisson denoising, and motion estimation in zoomed regions of cortex, hippocampus, and thalamus in Neuropixels 2.0 data. This is a screenshot of supplementary video for figure 4 in main text showing a frame that corresponds to the hippocampal zoomed region. In each frame top left panel shows the sparse localization image of detected spikes and their spatial positions after having been projected along the depth (y) axis. Top right panel is analogous to top left panel but shows the projection along the horizontal axis (x). Middle panels show the localization images after having gone through Poisson denoising [9]. Bottom panels show the Poisson denoised images after motion estimation and registration. Note that in all anatomical regions, the motion has been mitigated in the registered panel.

### 6.3 Supplementary video for figure 5 in main text (Web link: video-figure-5.mp4)

This video compares the localization images and the registration performance using all three compared methods (ours, Hurwitz et al. and CoM) in both Neuropixels 1.0 and 2.0 datasets. Screenshot of this video is shown in figure 10.

**Figure 10:**
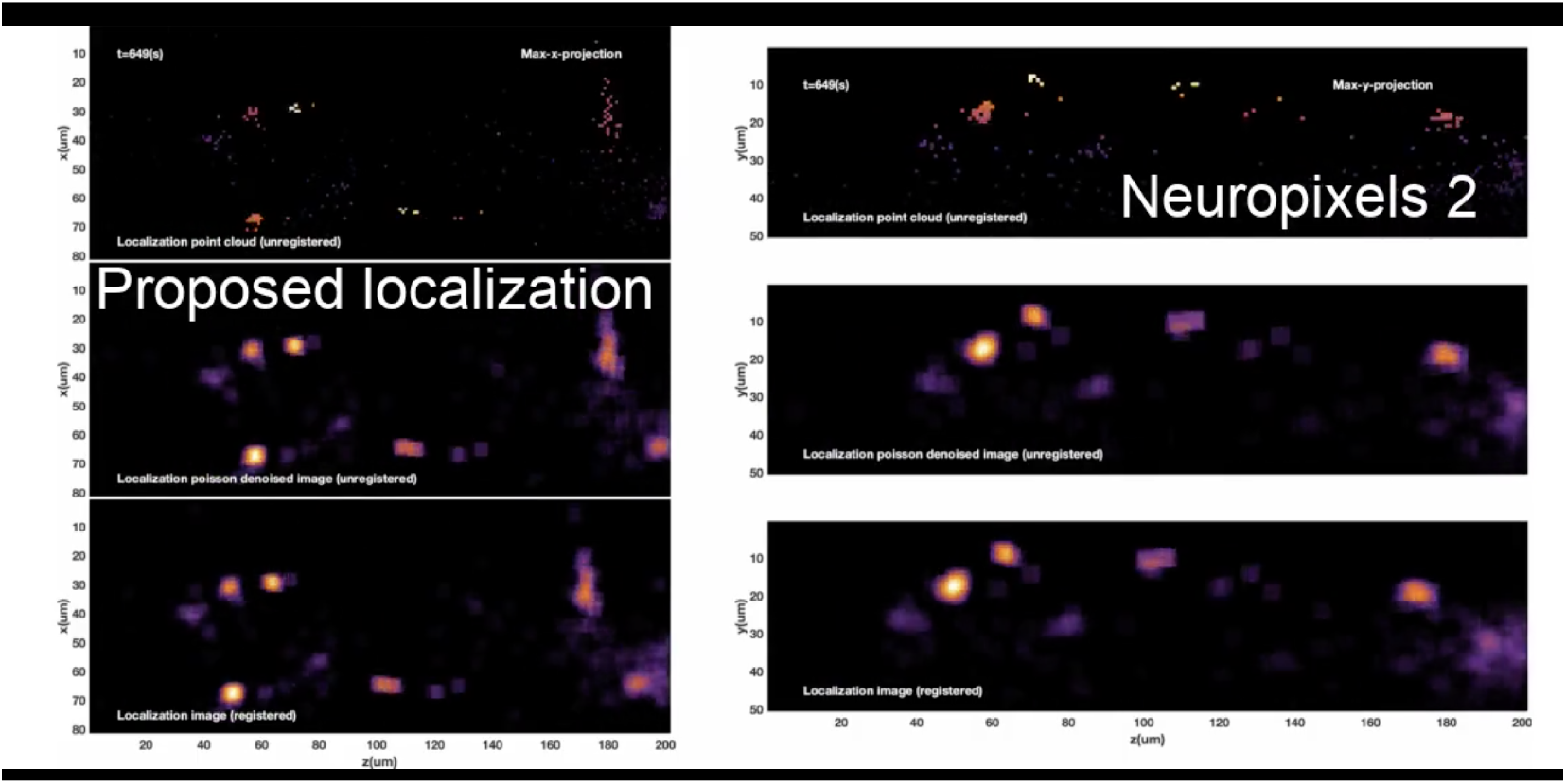
Video representation of Poisson denoising, and motion estimation using compared localization methods in both Neuropixels 1.0 and 2.0 data. This is a screenshot of supplementary video for figure 5 in main text showing a frame that corresponds to the cortical zoomed region using localization and registration guided by our technique in Neuropixels 2.0 data. Subsequent frames show comparisons with Hurwitz et al. and CoM in both Neuropixels 1.0 and 2.0 data. In each frame top left panel shows the sparse localization image of detected spikes and their spatial positions after having been projected along the depth (y) axis. Top right panel is analogous to top left panel but shows the projection along the horizontal axis (x). Middle panels show the localization images after having gone through Poisson denoising [9]. Bottom panels show the Poisson denoised images after motion estimation and registration. Note that the periodic horizontal motion visible in the middle (unregistered) panels is mitigated in the bottom panels after registration, demonstrating that the estimated motion corresponds well to the underlying motion. Also note that our proposed method yields a more stabilized registration. Since the CoM method compresses localization to a narrow strip, features that could be used to guide a fine resolution registration are lost, causing an underestimation of motion.

## 7 Datasets

For reproducibility of our experiments, datasets used in our paper can be found here: https://github.com/flatironinstitute/neuropixels-data-sep-2020/blob/master/doc/cortexlab1.md

